# Peptide location fingerprinting reveals modification-associated biomarkers of ageing in human tissue proteomes

**DOI:** 10.1101/2020.09.14.296020

**Authors:** Matiss Ozols, Alexander Eckersley, Kieran T Mellody, Venkatesh Mallikarjun, Stacey Warwood, Ronan O’Cualain, David Knight, Rachel EB Watson, Christopher EM Griffiths, Joe Swift, Michael J Sherratt

## Abstract

Although dysfunctional protein homeostasis (proteostasis) is a key factor in many age-related diseases, the untargeted identification of structural modifications in proteins remains challenging. Peptide location fingerprinting is a proteomic analysis technique capable of identifying structural modification-associated differences in mass spectrometry (MS) datasets of complex biological samples. A new webtool (Manchester Peptide Location Fingerprinter), applied to photoaged and intrinsically aged skin proteomes, can relatively quantify peptides (spectral counting) and map statistically significant differences to regions within protein structures. New photoageing biomarkers were identified in multiple proteins including matrix components (collagens and proteoglycans), oxidation and protease modulators (peroxiredoxins and SERPINs) and cytoskeletal proteins (keratins). Crucially, for many extracellular biomarkers, structural modification-associated differences were not correlated with relative abundance (by ion intensity). By applying peptide location fingerprinting to published MS datasets, (identifying biomarkers including collagen V and versican in ageing tendon) we demonstrate the potential of the MPLF webtool to discover novel biomarkers.

## Introduction

Loss of proteostasis (the ability to regulate the proteome) is a key feature of many age-associated diseases (1). In human skin, for example, ageing induces remodelling of the architecture and abundance of key structural proteins (such as fibrillar collagen and elastic fibres) which impact on appearance and function. This loss of proteostasis is particularly evident in sites exposed to ultraviolet radiation (UVR) (2). The photoageing process in these sites differentially affects the outer keratinocyte-rich epidermis and the lower extracellular matrix (ECM)-rich dermis (3, 4). Within the dermis, some ECM macromolecular assemblies, such as collagen 1 and elastic fibres are known to be long-lived with biological half-lives spanning years and decades (5, 6). These abundant structural proteins are thought to accumulate photoageing-induced damage over time (7) as a consequence of chronic ultraviolet radiation (UVR) exposure (predominantly UVA) (8–10), UVR-induced reactive oxygen species (ROS) (11, 12) and UVR-induced elevated protease activity (13). It is not known, however whether protein modifications as a consequence of ageing is a common feature of ECM proteins in the complex dermal matrisome. In contrast to the dermis, epidermal keratinocytes and biomolecules are further exposed to higher energy UVB as well as to lower-energy UVA, leading to the progressive accumulation of DNA damage (14) and a time-dependent modulation in gene expression and potential loss of function (15). As a consequence of higher protein turnover, it is likely that damage manifests primarily in the epidermis as changes in the ability of cells to synthesise functional proteins (16). Due to the biological complexity of skin and these varied mechanisms of damage, the photoageing process creates a spectrum of modifications to proteins which are challenging to track and distinguish using a single methodology. It is therefore necessary to develop novel methods of analysis for the detection of biomarkers characteristic of photoaged skin.

Label-free proteomic liquid chromatography tandem mass spectrometry (LC-MS/MS) is a powerful analytical technique used traditionally for the identification and relative quantification of proteins within complex mixtures, often extracted from whole tissues. The technique has enabled the identification of disease and ageing biomarkers based on their relative abundance (17, 18). Although this approach may successfully identify disease and age-related changes in protein abundance and deposition in ECM-rich tissues, it is poorly suited to the detection of abundance-independent, damage-associated protein modification in complex protein mixtures.

In order to address the need for an approach capable of identifying proteins with photoageing specific-modifications, we have developed the publicly available Manchester Peptide Location Fingerprinter (MPLF) webtool which can detect changes in peptide yield along protein structure (10, 19). MPLF is an LC-MS/MS proteomic analysis tool which implements peptide location fingerprinting: tryptic peptide spectral counts are mapped to protein regions (specified by the user, or from the UniProt database), relatively quantified per region and statistically tested between groups (**Fig. 1**). This enables the characterisation of regional differences as a consequence of structure-related modifications.

**Figure 1.**
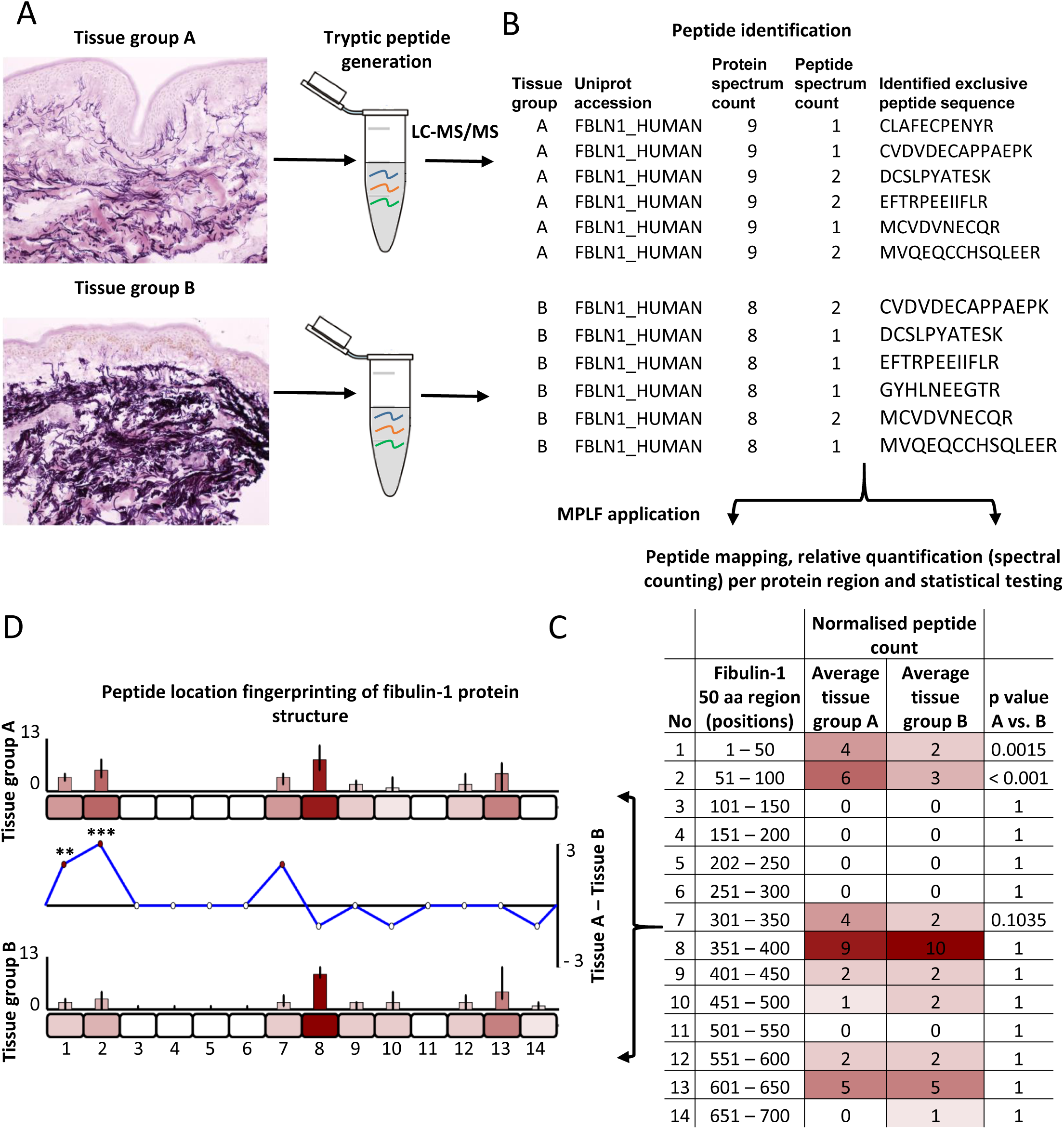
Identification of regional peptide yield differences within proteins using peptide location fingerprinting. As standard in proteomic MS, proteins are first extracted from tissues of interest using study-specific protocols and trypsin digested **(A)**. Post LC-MS/MS, exclusive (protein-specific) tryptic peptide sequences are identified and counted (spectral counted, i.e. peptide spectrum matches; PSMs) per tissue sample. Fibulin-1 is used throughout this figure as an exemplar **(B)**. Peptide count reports are uploaded to the MPLF webtool **(C)** where protein structures are divided either into user-defined amino acid (aa) step sizes (e.g. 50 aa shown here) or into step sizes corresponding to domains, repeats or regions pre-defined by the UniProt database. Peptides and their counts are then mapped to their protein regions and summed. Sample-specific regional counts are then normalised across the experiment based on the median protein spectrum count, averaged per tissue group and statistically compared (Bonferroni-corrected repeated measures ANOVA). This analysis is then visualised using representative amino acid-scale schematics of each protein **(D)**. Average peptide counts are heat mapped to their corresponding region for comparison between groups (bar graphs). Regional, average peptide counts in one tissue group are subtracted from the counts of the other to show regional differences in peptide yield (line graph) with statistical significances indicated (** ≤ 0.01, *** ≤ 0.001).

Using peptide location fingerprinting, we have previously shown that: i) long-lived macromolecular ECM assemblies exhibit inter-tissue structural diversity (19) and; ii) *in vitro* UVR-induced damage modifications to protein tertiary/quaternary structures can be detected in cell culture-derived suspensions of isolated ECM assemblies (fibrillin and collagen 6 microfibrils) and their associated receptors (integrins) (10). Here, we show that this same method can be used as a surveying tool capable of detecting statistically significant, photoageing-specific, structure modification-related differences in proteins within human epidermal and dermal proteomes. In addition to this, the MPLF webtool was further used to analyse a previously published human ageing tendon dataset (20) to demonstrate the webtool’s wider applicability in evaluating existing LC-MS/MS data. This enables the identification of potentially novel biomarkers and protein classes which are independent from protein abundance when compared to traditional relative quantification by peak area ion intensity.

## Results and Discussion

### Peptide location fingerprinting reveals regional fluctuations in peptide yield within the structures of proteins from photoaged skin

Prior to LC-MS/MS we confirmed that forearm skin in our donor cohort was consistently and severely photoaged compared with buttock skin. In comparison to buttock skin sections from aged donors, matched extensor forearm skin exhibited clear hallmarks of photoageing (**Fig. S1**) including epidermal thinning and disruption of elastic fibre architecture including solar elastosis (21, 22). Quantification of solar elastosis (% elastic fibre coverage) indicated that forearm skin had significantly higher abundance of elastin in comparison to matched buttock skin (**Fig. S2**).

As the epidermis and dermis contain distinct molecular and cellular populations, we separated these two skin layers and analysed them independently with LC-MS/MS. A total of 51,895 protein-specific tryptic peptide sequences corresponding to 975 proteins (**Table S1**) were detected across all dermal samples and 45,868 sequences corresponding to 836 proteins across epidermal samples (**Table S2**). Principal component analysis (PCA) of peptide spectral counts showed clear separation of extensor forearm and buttock data into distinct clusters (**Fig. S3**) highlighting the global difference between photoaged and intrinsically aged skin. Peptide sequences were then regionally mapped to their respective proteins, relatively quantified for forearm and buttock and statistically compared using the MPLF webtool. A full workflow describing the application of peptide location fingerprinting using the MPLF webtool to our skin datasets for photoageing biomarker identification is shown in **Fig. 2**. Online access to webtool will be provided upon publication in a scientific journal.

**Figure 2.**
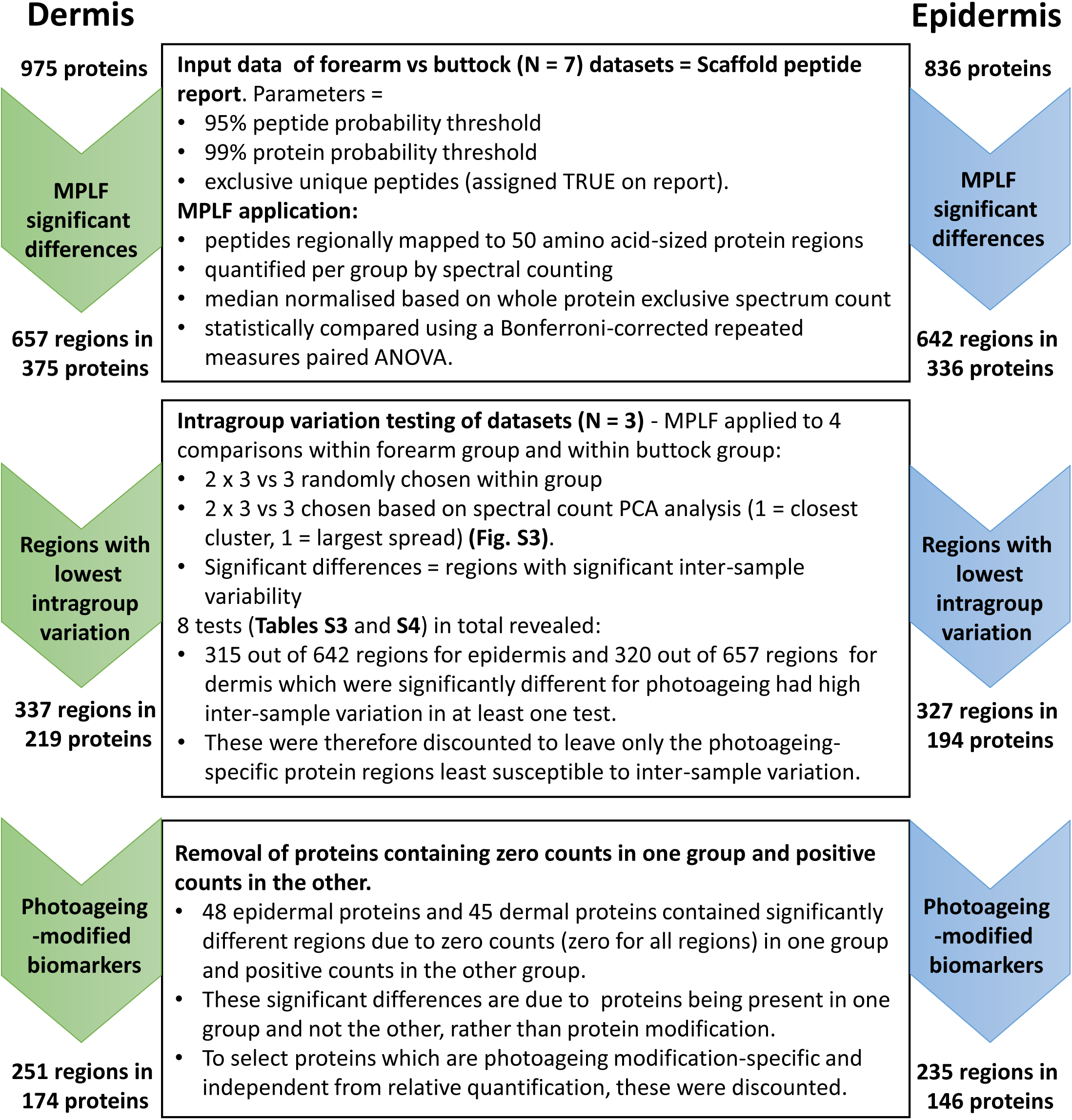
Identification of photoageing-modified biomarkers in human skin by filtering significant regional differences detected using the MPLF webtool. The MPLF webtool was applied to dermal and epidermal datasets to identify 50 aa-sized protein regions with significant differences in peptide yield between forearm and buttock samples. Once identified, (657 for dermis, 642 for epidermis) significantly different regions were tested for intragroup variation (forearm group and buttock group; **Tables S3** and S4). Regions which exhibited significant intragroup variation were discounted to leave only those which were photoageing-specific (337 for dermis, 327 for epidermis). Lastly, proteins which were present in one group (forearm or buttock) but not the other were also discounted. The remaining regions (251 for dermis, 235 for epidermis) and their respective proteins were considered as possible structural modification-specific biomarkers of photoageing.

Using the MPLF webtool, peptide location fingerprinting identified a total of 657 protein regions in dermis and 642 regions in epidermis (corresponding to 375 and 336 proteins respectively) which had significant fluctuations in peptide yield between photoaged forearm and intrinsically aged buttock samples (for standardisation, all regions were 50 amino acids in size; **Fig. 2**). Of the protein regions displaying significant differences of peptide counts between forearm and buttock samples, 337 for dermis and 327 for epidermis were least susceptible to intragroup variation (**Tables S3** and S4) and therefore most photoageing-specific (these corresponded to 219 proteins in dermis and 194 proteins in epidermis). Lastly, to identify modification-specific regional differences that are most independent from protein presence, proteins which were present in either forearm or buttock groups but not the other were also discounted. This left a total of 251 regions within 174 proteins in dermis (**Table S5**) and 235 regions within 146 proteins in epidermis (**Table S6**) to be considered as biomarkers susceptible to photoageing-related modifications (**Fig. 2**).

Of the protein biomarkers identified with local molecular differences using peptide location fingerprinting, eight exemplar proteins that play key functional roles in skin are displayed in **Fig. 3** (interleaved graphs: **Fig. S4**). Collagen 6 alpha-3, fibulin-1, biglycan and galectin-7 from the dermis and keratins (K)-2 and -10, desmoplakin and heat shock protein (HSP) 70 from the epidermis all had significant regional differences in peptide yields along their structures. This indicated that these proteins were structurally modified as a consequence of photoageing.

**Figure 3.**
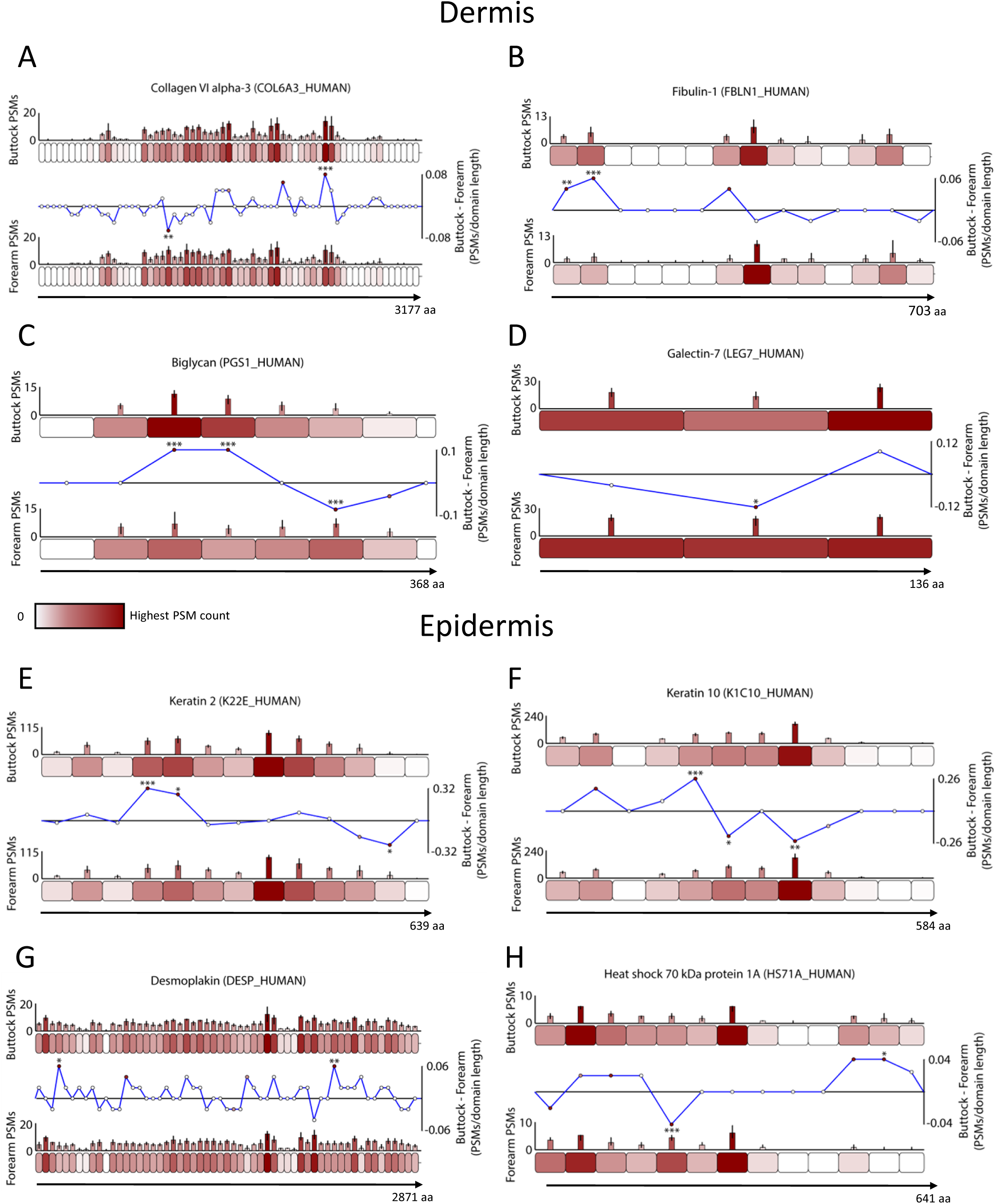
Exemplar skin biomarkers exhibiting photoageing-specific structural modifications. Proteins were segmented into 50 aa-sized step regions with average peptide counts (PSMs; N = 7) heat mapped to each step and compared between forearm and buttock (bar graphs = average PSMs, error bars = SD). Average peptide counts corresponding to each forearm protein step were subtracted from the counts of their corresponding buttock protein step and divided by the amino acid sequence length of that step to reveal regional differences in peptide yield (line graphs). Multiple protein structures within the dermis **(A - D)** and epidermis **(E - H)** exhibited statistically significant regional differences in the peptide yield (* ≤ 0.05, ** ≤ 0.01, *** ≤ 0.001; Bonferroni-corrected repeated measures paired ANOVA) between forearm and buttock. In the dermis, one region in the N-terminal half of collagen 6 alpha-3 had significantly lower peptide counts in buttock samples than in forearm, whereas another region in the C-terminal half of the protein had significantly higher **(A)**. Two N-terminal regions within fibulin-1 had significantly higher peptide counts in buttock than in forearm **(B).** Biglycan had two regions near the central portion of the protein with significantly higher peptide counts in buttock than in forearm and another region on the C-terminal side which had significantly lower **(C).** A central protein region in galectin 7 yielded significantly lower peptides in buttock samples than in forearm **(D)**. In epidermis, two regions in K2 **(E)** and one in K10 **(F)** (on the N-terminal sides of both proteins) had significantly higher peptide counts in buttock than in forearm whereas one region on the C-terminal end of K2 and two near the protein centre of K10 had significantly lower counts in buttock than forearm. In desmoplakin **(G)** one N-terminal region and another near the C-terminus yielded significantly higher peptides in buttock than in forearm. Lastly, heat shock protein 7 exhibited one N-terminal sided region with significantly lower peptide counts in buttock than in forearm and another C-terminal region with significantly higher **(H)**.

The collagen 6 microfibril is a structural ECM assembly comprised predominantly of three alpha chains with alpha-3 being the largest (23). Although previous histological analysis of photoaged skin showed no gross changes to the distribution of the collagen 6 network (24), we recently demonstrated using peptide location fingerprinting that the alpha-3 chain was structurally susceptible to physiological doses of UVR *in vitro* (10). As these ECM assemblies are markedly long-lived, here we show evidence for the first time that the alpha-3 chain is susceptible to photoageing-dependant modifications *in vivo*. Fibulin- 1 is an ECM-associated protein (25, 26) in the dermis which influences cell activity by modulating integrin interactions (27). Although skin photoageing has been shown to affect the co-localisation of fibulins-2 and -5 on elastic fibres (25, 28), we demonstrate that the structure of fibulin-1 may also be affected. Biglycan is an ECM-associated proteoglycan (29) capable of transforming growth factor-β (TGFβ) regulation (30) whose presence is reduced in photoaged forearm dermis compared to intrinsically aged buttock (31). We reveal that the biglycan present in photoaged forearm is also subjected to damage modification by photoageing which may impair functionality. Galectin-7 plays a key role in epidermal cell migration and wound re-epithelialisation (32). Although reductions in epidermal galectin-7 expression was shown recently in intrinsically aged skin (33), our study demonstrates that dermal galectin-7 is additionally susceptible to modifications as a consequence of photoageing.

Within the epidermis, both K2 and K10 had evidence of being structurally modified in photoageing. Both are heavily involved in keratinocyte differentiation and cornification (34). Although the upregulation of several epidermal keratins (35) has been shown in response to acute UVR exposure, including K10 (36), differences to K2 and K10 as a consequence of chronic UVR exposure have not to our knowledge been previously been demonstrated *in vivo*. The desmosomal protein desmoplakin is required for epidermal cell adhesion (37). To our knowledge, this is the first instance of a photoageing-specific difference to an epidermal desmosomal protein. HSP70s are molecular chaperones and crucial regulators of cell proteostasis, a process that becomes increasingly unbalanced during ageing (38). In mice, HSP70 overexpression was shown to suppress UVR-induced wrinkle formation highlighting its therapeutic potential (39). However, this is the first demonstration of photoageing-dependant alterations to HSP70.

Through a variety of mechanisms initiated by chronic UVR exposure (direct photochemistry, induction of ROS and aberrant protease expression and activity), photoageing introduces a spectrum of protein modifications which are very challenging to track. Peptide location fingerprinting successfully enabled the identification of local molecular differences within protein regions revealing a profile of proteins with photoageing-specific modifications. These may impact the functionality of skin, influencing multiple mechanisms of damage and, ultimately, skin homeostasis. For this reason, it is important to next consider these gross changes in infrastructure with a more holistic approach using classification and pathway analysis.

### Classification and pathway analysis of skin proteins with photoageing modifications reveals ECM, cytoskeletal proteins and metabolic enzymes as classes most affected

Classification analysis of proteins with significant differences in regional peptide yield between forearm and buttock highlighted components of the ECM as the most affected protein class in the dermis (**Fig 4A**, blue pie chart). Common to both dermis and epidermis, cytoskeletal proteins (grey pie chart; top in epidermis) and metabolite interconversion enzymes were also among the top four classes most affected by photoageing, according to peptide location fingerprinting. Protease activity modulators (yellow pie chart) were confined to the dermis and translational (**Fig. 4B**, brown pie chart) and DNA- / RNA-binding proteins (dark blue pie chart) were classes exclusive to the epidermis. Crucially, this analysis identifies the potential involvement of several new protein families in the photoageing process, such as peroxiredoxins, serpins, ribosomal proteins and heterogeneous nuclear ribonucleoproteins (hnRNPs), as well as collagens, laminins, proteoglycans and keratins with previously unappreciated roles in photoageing biology.

**Figure 4.**
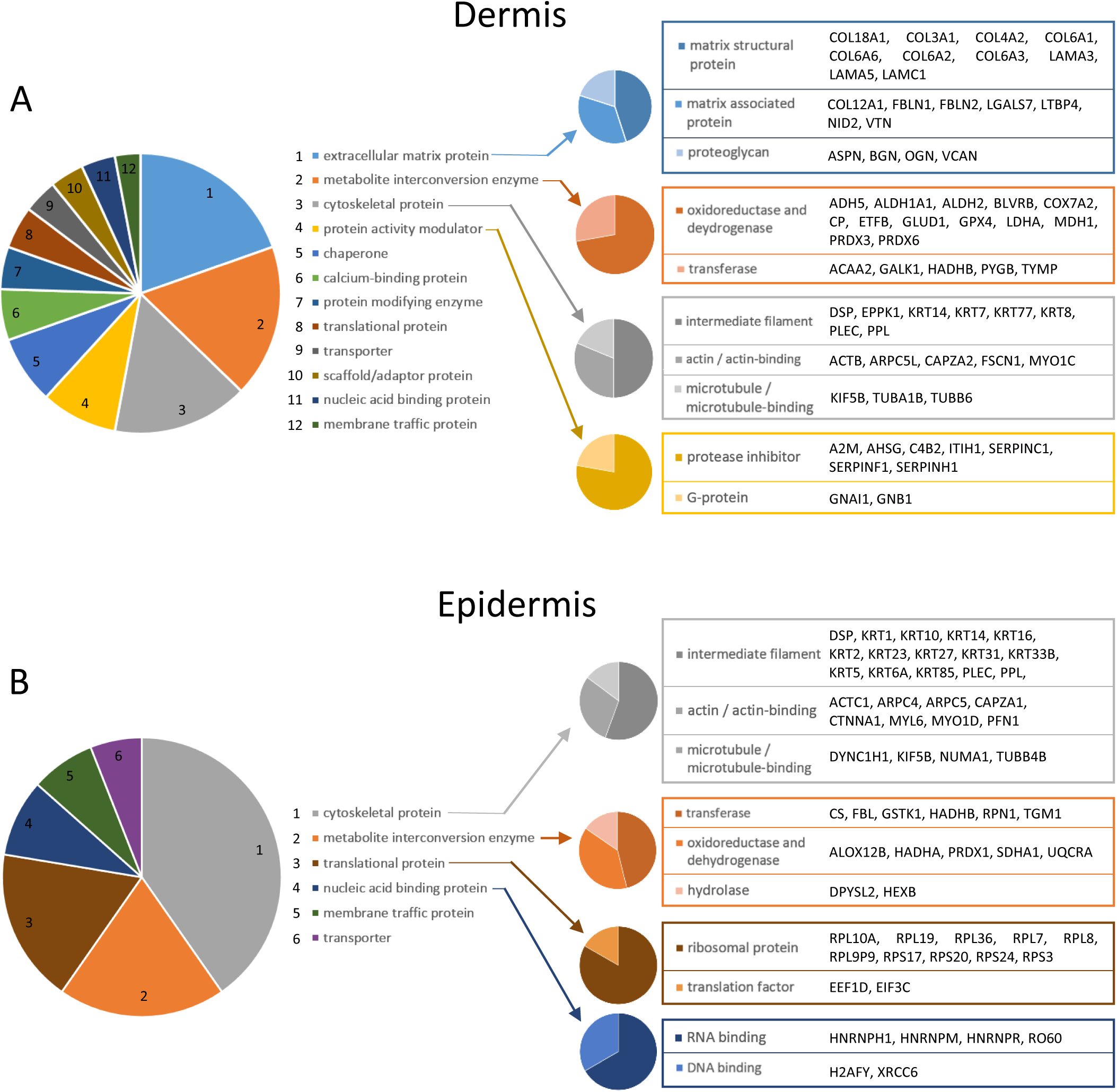
Classification of modification-specific protein biomarkers into functional groups reveals ECM components, ROS- and protease-modifying enzymes, cytoskeletal proteins and ribosomal proteins as the main classes in skin most affected by the photoageing process. Biomarker proteins with photoageing-specific modifications were categorised into protein classes (PANTHER classification system; large multi-coloured pie charts: clockwise rankings with top rank at 12:00; only classes with two or more proteins are represented). The top four classes for dermis **(A)** were: ECM proteins (blue slice) which can be further categorised (blue pie chart) into structural matrix proteins, matrix-associated proteins and proteoglycans; metabolite interconversion enzymes (orange slice) which can be categorised (orange pie chart) into oxidoreductase and transferase enzymes; cytoskeletal proteins (grey slice) which can be categorised (grey pie chart) into intermediate filament-, actin-, and microtubule-associated proteins; and protein activity modulators (yellow slice) which can be categorised (yellow pie chart) into protease inhibitors and G-proteins. As for the dermis, cytoskeletal proteins and metabolite interconversion enzymes were also in the top four classes for the epidermis **(B)** in addition to translational proteins (brown slice) which can be categorised (brown pie chart) into ribosomal proteins and translational factors and nucleic acid binding proteins (dark blue slice) which can be categorised (dark blue pie chart) into RNA- and DNA-binding proteins. Protein identities contained within those categories are also listed on the right.

The dermis is comprised primarily of long-lived structural ECM proteins. Some of the structural ECM biomarkers identified by peptide location fingerprinting such as collagen family members have half-lives measured in decades (6, 40). Therefore, many of these assemblies are thought to accumulate damage as a result of chronic UVR exposure, leading to changes in molecular abundance and function, ultimately contributing to the photoageing phenotype (7, 41, 42). The majority of evidence for this is based on histological differences in elastic fibre (such as by fibrillin-1 and elastin) and collagen 1 architecture and protein abundance. Although peptide location fingerprinting did identify significant differences in the protein regions within fibrillin-1 and collagen 1 alpha chains (**Table S5**), the high intragroup variation of these regions between samples (**Table S3**) meant that it was difficult to attribute differences to either photoageing or individual variability. Due to the stringency of our methods, these were therefore disregarded as modification-dependant biomarkers of photoageing. For proteins such as these, with higher inter-individual variability, more samples may be required to identify photoageing-specific regional changes. Regardless, peptide location fingerprinting revealed numerous, potentially novel ECM biomarkers meriting further study (**Fig. 4A**), whose regions were not susceptible to individual variation according to our methods. These include alpha chains of collagens 3, 4 and 12, elastic fibre-associated proteins such as fibulins-1, -2 and LTBP4, basement membrane nidogen-2 and laminins A3, A5 and C1, and lastly the proteoglycans asporin, biglycan, mimecan and versican. These novel biomarkers were also highlighted within protein-protein interaction networks using STRING analysis, which showed a cluster of interactions between these proteins (collagens in particular; **Fig. S5**, blue dashed line). The identification of these ECM protein super-families strengthens the hypothesis that long-lived macromolecular assemblies accumulate damage over time which may be independent from protein presence or abundance.

In addition to versican, which has previously been identified as a dermal biomarker of skin photoageing (43), asporin, lumican and biglycan all bind to fibrillar collagens (44–46) and are capable of modifying their assembly and function (47). Similarly, versican binds to fibrillin microfibrils and link these major elastic fibre component to connective tissue networks (48). As such, these proteoglycans function to build and network major structural ECM components. We hypothesise therefore, that their functional decline may lead to the degeneration of matrix architecture seen in photoaged forearm (49).

The ROS-modulating (50) peroxiredoxins (PRDXs)-3 and -6, identified by peptide location fingerprinting (**Fig. 4**), have not previously been reported as affected by photoageing. Photoexposed tissue is commonly associated with an increased oxidative environment (51) through the photodynamic

formation of ROS (11, 52). The functional decline of these antioxidants may play an active part in the progression of photoageing.

Of the protease modulators, three serine protease inhibitors (SERPINs) were also newly identified (**Fig. 4A**). As well as the inhibition of serine proteases (53) some, such as SERPINA1, also inhibit collagenase matrix metalloproteinase (MMP) 2 and the gelatinase MMP9 preventing the degradation of collagen 1 (54), which is reduced in photoaged dermis (41). The structure-associated modifications identified within three members of the SERPIN superfamily may be indicative of their impaired function and therefore synonymous with a global dysregulation of dermal protease activity.

Within both dermal and epidermal proteomes, intracellular metabolite interconversion enzymes and structural cytoskeletal proteins were two of the main classes with significant regional differences between photoaged forearm and buttock (**Fig. 4**). These new observations suggest that the photoageing process may be having a global effect on cell metabolism and structure. Although the acceleration of cumulative DNA damage as a result of chronic UVR exposure may not necessarily lead to tumorigenesis, it is still thought to have a profound impact on keratinocyte function. This includes disordered maturation, a loss of polarity (15) and a decrease in the capacity for differentiation and proliferation *in vivo* (55). It is likely that our identification of a large number of proteins fundamental to cell metabolism and structure may reflect these large-scale changes in cell function and behaviour which are characteristic of epidermal chronic sun exposure.

Of particular interest, within the epidermal proteome (in addition to K2 and K10, **Fig. 3E, F**), a number of other keratins were also identified as modified by photoageing (**Fig. 4B**, grey pie chart). This is also highlighted by STRING analysis which shows a cluster of interactions between all twelve keratin members identified (**Fig. S6**, dashed blue line). These keratins play a variety of roles within the epidermis from cornification (56) and barrier function integrity (57) in the superficial layers to keratinocyte differentiation and proliferation in deeper layers (58). The revelation of a global change within the keratin superfamily by peptide location fingerprinting is indicative of a wider functional decline of these crucial processes.

Peptide location fingerprinting revealed that several epidermal proteins affected by photoageing were ribosomal (**Fig 4B**, brown pie chart). STRING analysis also demonstrates a cluster of interactions between all ten ribosomal protein members identified (**Fig. S6**, dashed black line). Structural changes in these proteins may result in the decline of ribosome functionality within resident epidermal cells. The deterioration of ribosome presence (59) and function (60) during ageing has been shown previously in mice. In addition, it is well established that photoageing grossly affects gene expression in human keratinocytes (16, 61). A decline in ribosome function and in RNA-binding proteins (**Fig 4B**, blue pie chart) would lead to impairment of the cell’s translational machinery impacting globally on keratinocyte gene expression.

Peptide location fingerprinting allows the assessment of protein modifications both on a molecular scale by visualising significant differences in peptide yield along a protein’s structure (**Fig. 3**) and on a proteomic scale through global analyses of identified biomarkers (**Fig. 4**). Together, it reveals potentially novel mechanisms and/or perturbed pathways following chronic sun exposure which merit future functional validation and study.

### Peptide location fingerprinting reveals biomarkers of the photoageing process which are not identified by conventional relative quantification of protein abundance

In addition to regional protease susceptibility using peptide location fingerprinting, protein abundance was also relatively quantified within the same label-free LC-MS/MS datasets by peak area ion intensity. This was performed in order to identify abundance-dependant protein differences due to photoageing and reveal biomarkers unique to both methodological approaches. A total of 635 dermal proteins and 926 epidermal proteins were significantly different in relative abundance (**Fig. S7**) between photoaged forearm and intrinsically aged buttock according to peak area intensity (principal component analyses indicated clear data separations, **Fig. S8**; full list of proteins - dermis: **Table S7**, epidermis: **Table S8**). As seen for peptide location fingerprinting, classification analysis also identified cytoskeletal proteins, metabolite interconversion enzymes and translational proteins as the top three classes whose relative abundance was affected in forearm epidermis (**Fig. S9**). These may be linked to the profound changes to keratinocyte morphology and gene expression as a result of the photoageing process (62). Counterintuitively, the main protein classes affected in dermis were intracellular rather than ECM-associated. A lower presence of fibroblasts in intrinsically aged skin (buttock) (63) coupled to the higher biosynthetic activity of fibroblasts and an increased presence of inflammatory cells in photoaged skin (forearm) (64) may explain these differences in intracellular protein abundance.

Two new potential photoageing biomarkers TIMP3 (metalloprotease inhibitor 3) in dermis and RPL36 (ribosomal protein L36) in epidermis, identified as significantly different in relative abundance between forearm and buttock skin by LC-MS/MS peak area intensity, were successfully experimentally validated with Western blotting (**Fig S10**). Western blotting on the same dermis and epidermis protein extracts corroborated a significant increase in TIMP3, and decrease in RPL36, in photoaged forearm compared to intrinsically aged buttock. Furthermore, normalised relative quantification of TIMP3 and human serum albumin (loading control selected based on non-significance) by Western blotting matched well with that of LC-MS/MS peak area relative quantification on a sample by sample basis (**Fig S11**). Although all four TIMPs have been shown to inhibit all 26 MMPs (65), TIMP3 is also capable of inhibiting more members of the a disintegrin and metalloproteinase proteins (ADAMs) and ADAMs with thrombospondin motifs (ADAMTSs) than its other family members (66). One proposed mechanism of photoageing is that chronic exposure of UVR to fibroblasts and keratinocytes can lead to the upregulation of MMPs in skin (13, 67). The elevated presence of TIMP3 in photoaged forearm dermis compared to intrinsically aged buttock is perhaps indicative of a heightened proteolytic environment and an attempt to mitigate UVR-induced remodelling.

Peptide location fingerprinting uniquely identified 120 protein biomarkers in the dermis (**Fig. 5A**; **Table S9**) and 71 biomarkers in the epidermis (**Fig. 5B**; **Table S10**), which were modified as a consequence of photoageing but did not differ significantly in relative abundance according to peak area quantification. This demonstrates that changes in protein abundance do not necessarily correlate with changes related to protein modification, particularly in ECM-rich tissues. Of these, thirteen were ECM-associated proteins including four collagens, two elastic fibre-associated proteins (LTBP4 and nidogen- 2), three laminin chains, nidogen-2 and the proteoglycan versican. This suggests that peptide location fingerprinting is capable of identifying long-lived proteins (5, 6, 68) subjected to accumulated chronic damage (7), whose abundance may not be impacted by the photoageing process. This is also corroborated by protein classification analysis which ranked ECM proteins as the top most affected classes in dermis for structural modifications (**Fig. 4**), but not for changes in relative abundance (**Fig S9**). In epidermis, peptide location fingerprinting also successfully identified photoageing modifications in numerous keratins, serpins and ribosomal proteins which did not significantly differ in abundance.

**Figure 5.**
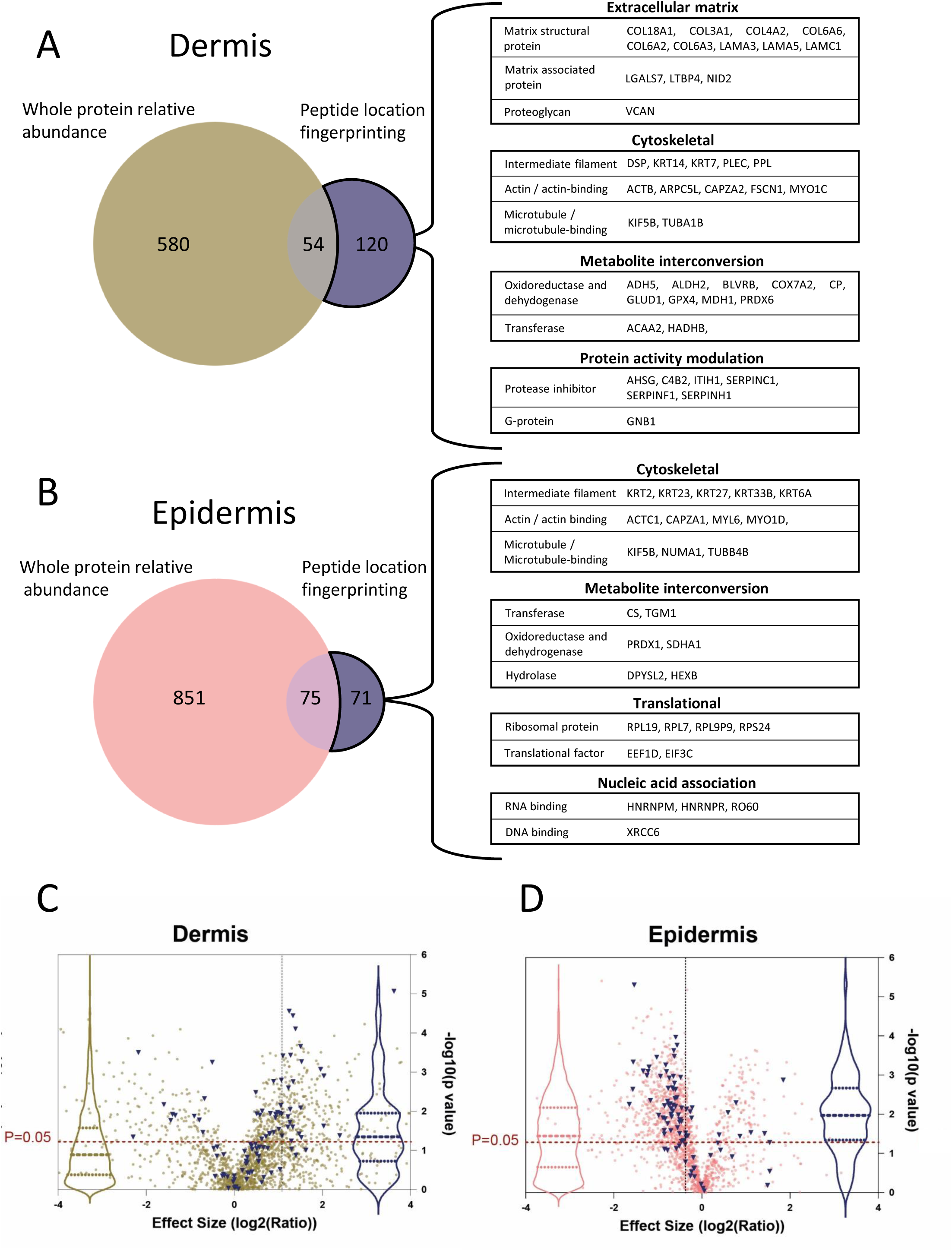
Peptide location fingerprinting identifies unique modification-specific skin biomarkers of photoageing which were not identified by whole protein relative quantification. Venn diagrams compare the number photoageing-modified protein biomarkers identified by peptide location fingerprinting with the number proteins identified with significant differences in whole protein relative abundance by peak area quantification. Several proteins were uniquely identified by peptide location fingerprinting as photoageing-specific biomarkers in both dermis **(A)** and epidermis **(B)** including multiple ECM proteins and protein activity modulators for dermis, translational and nucleic acid-associated proteins for epidermis and cytoskeletal proteins and metabolite interconversion enzymes for both sub-tissues. Protein identities with significant modifications in structure correlated less strongly with significant differences in protein abundance in the dermis **(C)** than in the epidermis **(D)**. Proteins with photoageing-specific structural modifications were identified and labelled blue on relative protein abundance volcano plots from **Fig S7**. Violin plots (dashed lines = median and IQR) of the labelled points (blue) were superimposed to show correlation between proteins with modification-associated differences and protein abundance differences (violin plots of all data points also shown in pink and gold). For the epidermis, a large proportion of proteins which have structural modifications (blue points and blue violin plot) are positioned above the p = 0.05 line, indicating that these were also significantly different in relative abundance between photoaged forearm and intrinsically aged buttock. This suggests that for epidermis, structure-related differences appear to correlate strongly with differences in abundance (median p-value = 0.01). For the dermis however, a smaller proportion of proteins identified with modification-associated differences are positioned above the p = 0.05 line suggesting that these differences appear to correlate less with abundance (median p-value = 0.04).

To test whether differences in structural modification in proteins may correlate with differences in relative abundance in epidermis and dermis, p-values of relative abundance differences (measured by peak area) of proteins identified with modifications (measured by peptide location fingerprinting) were compared between epidermis and dermis (**Fig. 5 C, D**). The proportion of proteins with both significant modifications and also significant differences in relative abundance was higher in epidermis than in the dermis. Since the dermis consists primarily of longer-lived structural ECM whereas the epidermis is predominantly cellular, this observation may be reflective of the higher protein turnover in the cellular epidermis compared to the relatively acellular dermis. This is a further demonstration of the potential of peptide location fingerprinting in elucidating biomarkers of chronic disease, particularly in ECM-rich connective tissue but also in cell-rich tissues.

### The MPLF webtool can be readily applied to existing LC-MS/MS datasets to reveal previously undiscovered disease biomarkers

To showcase that peptide location fingerprinting can be used as a surveying tool to reveal novel disease biomarkers by post hoc analysis of published LC-MS/MS datasets, the MPLF webtool was used to analyse an independent, publicly available label-free LC-MS/MS human tendon data (downloaded from the PRIDE repository; dataset identifier PXD006466 and 10.6019/PXD006466). Hakimi et al. (2017) recently published a quantitative proteomic comparison between aged (torn) and young human supraspinatus tendon (20). They revealed significant reductions in the multiple pericellular matrix components in aged, torn tendon compared to young uninjured tendon, such as collagens 1 and 6 and elastic fibre components fibrillin-1, microfibril-associated protein 5 (MFAP5), LTBP2 and fibulin-1.

Peptide location fingerprinting on this published dataset identified 54 protein regions with significant differences in tryptic peptide yield between aged torn tendon and young uninjured male tendon (**Table S11**) corresponding to 31 proteins with possible modifications to structure (**Fig. 6A**). Of these, lamin-A, human serum albumin, tenascin-X and cartilage intermediate layer protein-2 (CILP2) are displayed as exemplary proteins with significant regional differences in peptide yields along their structures (**Fig. 6 B – E**).

**Figure 6.**
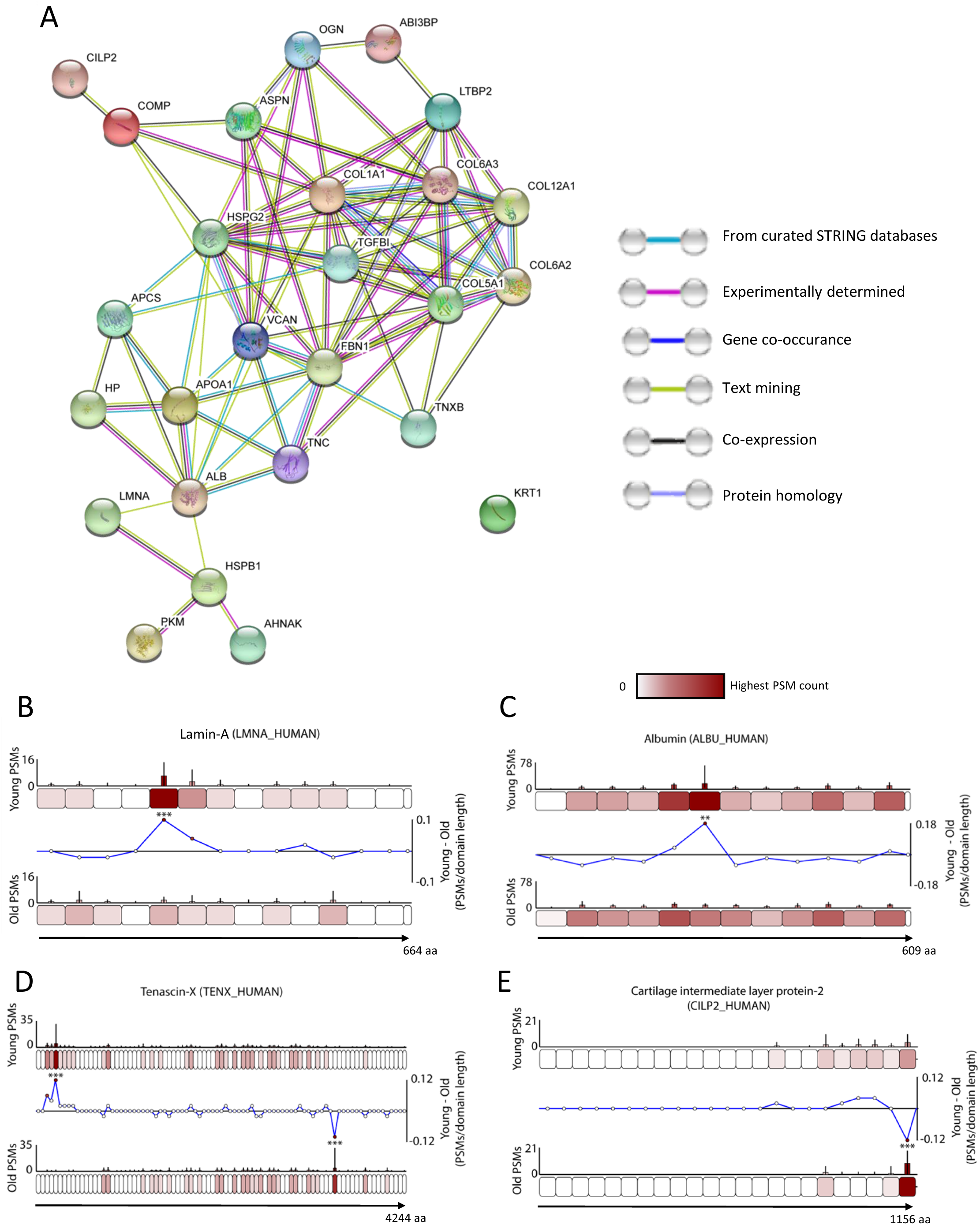
MPLF webtool analysis of publicly available ageing human tendon data demonstrates that peptide location fingerprinting can be applied post hoc to downloaded conventional label-free LC-MS/MS datasets to reveal unique structural modification disease biomarkers. Protein-protein interaction network (STRING; minimum required interaction score = 0.400) highlighted ECM and ECM-associated proteins as the main MPLF-identified tendon biomarkers of human ageing and injury **(A)**. Differences in peptide yield along the protein structure of four of these biomarkers identified are shown **(B – E)**. Similar to Fig. 2, proteins are segmented into 50 aa-sized step regions with average peptide counts (PSMs; N = 9) heat mapped to each step and compared between aged/torn and young groups (bar graphs = average PSMs, error bars = SD). Average peptide counts corresponding to each protein step in the aged/torn group were subtracted from the counts of their corresponding protein step in the young group and divided by the amino acid sequence length of that step to reveal regional differences in peptide yield (line graphs) with statistically significances between groups shown (** ≤ 0.01, *** ≤ 0.001; Bonferroni-corrected repeated measures unpaired ANOVA). Both lamin-A **(B)** and albumin **(C)** contained one region on their N-terminal sides with significantly higher peptide counts in young compared to aged samples. Tenascin-X contained one region near the N-terminus which yielded significantly higher peptides in young compared to aged and another on C-terminal side which yielded significantly lower peptide counts **(D)**. A C-terminal region of CILP2 also had significantly lower peptide counts in young compared to aged **(E)**.

In common with biomarkers identified by Hakimi et al. (2017) by relative quantification (20), peptide location fingerprinting also identified collagen 1 alpha-1, collagen 6 alpha 2, fibrillin-1, latent-transforming growth factor beta-binding protein 2 (LTBP2), complement and CILP as modification-specific markers of tendon damage. However, a number of newly identified biomarkers exclusive only to protein structural modifications were also identified. These were collagen V alpha-1, the proteoglycans versican, mimecan and asporin, human serum albumin, the nuclear intermediate filament protein lamin A, and the multifunctional cytokine TGFβ.

Through the use of the MPLF webtool, peptide location fingerprinting was successfully applied to a previously published human tendon dataset, revealing possible novel modification-specific protein biomarkers of tendon damage not previously identified (20). This highlights the wider impact of peptide location fingerprinting as a novel surveying tool capable of enhancing biomarker discovery in combination with conventional relative quantification LC-MS/MS methods.

## Conclusion

Peptide location fingerprinting of label-free LC-MS/MS datasets generated from photoaged forearm and intrinsically aged buttock skin enabled the identification of structure-associated modifications to proteins as a consequence of chronic sun exposure. As well as a biomarker discovery tool, we demonstrated that peptide location fingerprinting can be used to investigate local molecular differences in tryptic peptide yield within structural regions of proteins in complex, whole tissue lysates. Crucially, this approach also identified proteins which did not exhibit significant differences in relative abundance. As such, peptide location fingerprinting revealed novel biomarkers of photoageing which remain undetectable by conventional relative quantification, in particular of long-lived ECM proteins.

Through the use of the MPLF webtool, peptide location fingerprinting can be applied to any label-free LC-MS/MS dataset suitable for conventional data-dependant acquisition. As such, a combinational approach of both peptide location fingerprinting and relative protein quantification of protein abundance enables a more complete assessment of tissue proteostasis as a consequence of chronic disease and creates a powerful tool for biomarker discovery.

## Materials and Methods

### Human tissue and materials

Chemicals were sourced from Sigma-Aldrich Co. Ltd (Poole, UK) unless otherwise stated. Human skin was collected from photoaged donors (N = 7, mean age = 70 years, range = 62 – 79 years, 3 males; Fitzpatrick skin phototype I-III) with informed and written consent following approval from The University of Manchester Research Ethics Committee (ref: UREC 15464). Samples were bisected, one half snap frozen (for LC-MS/MS) and one half embedded in Optimal Cutting Temperature compound (OCT; CellPath; Powys, UK) and snap frozen (for histology).

### Tissue cryosectioning and Weigert’s staining

Tissue (buttock and forearm) was cryosectioned at a thickness of 5 µm (OTF cryostat; Bright Instruments, Bedfordshire, UK). Three serial sections were collected per slide and stained with Weigert’s resorcin fuchsin (Merck; Darmstadt, Germany). In brief, cryosections were fixed in 4% [w/v] paraformaldehyde (PFA) in PBS and then submerged in staining solution, each step for 10 minutes at room temperature. Stained sections were dehydrated in graded industrial methylated spirit (IMS) (70% [v/v], then twice with 100%), cleared in xylene (5 minutes per step at room temperature) and then permanently mounted (DPX).

### Imaging and solar elastosis quantification

Weigert’s stained skin sections were imaged using brightfield microscopy (BX53 microscope; Olympus Industrial, Southend-on-Sea, UK). Solar elastosis was quantified by measuring the percentage area of positively-stained elastic fibres using ImageJ (NIH; Bethesda, MA, USA). Percentage areas of elastic fibres were measured automatically by thresholding the images by brightness using “Threshold Colour”. All coloured pixels, corresponding to the purple stained elastic fibres were then converted to a single colour and all other pixels were coloured white. Elastic fibre abundances were measured within ImageJ as a percentage of coloured pixels within the areas. Statistical comparisons were performed using GraphPad Prism (GraphPad Software Incorporated; California, USA). The percentage area of elastic fibres was assessed for matched forearm and buttock skin groups with six individuals per group. Percentage area was measured in three images per section and three sections per individual, and then averaged. Elastic fibres were analysed between matched forearm and buttock groups using the Student’s paired t-test. Only differences of p ≤ 0.05 were considered significant.

### Sample preparation for mass spectrometry

Employing any LC-MS/MS analysis on skin dermis is challenging as networking ECM structures are tightly bonded, highly modified and covalently crosslinked. This, along with the hierarchical assembly of macromolecular ECM components (as seen in fibrillary collagens and elastic fibres) renders them highly insoluble (69). As such, an optimised protocol was devised which maximised peptide yield and protein discovery in dermis and epidermis by LC-MS/MS analysis.

Skin samples were defrosted at room temperature and incubated in 20 nM Ethylenediaminetetraacetic acid (EDTA) in PBS for 2 hr at 37°C to ensure dermal-epidermal separation. Biopsies were washed in PBS and the epidermis removed. Dermis was minced and incubated in 8 M urea buffer (8 M urea + 25 mM ammonium bicarbonate + 25 mM dithiothretol [DTT]) and homogenised using a bullet blender (Next Advance, New York, USA) at maximum speed. Dermal samples were centrifuged and supernatants diluted to 2 M urea with AB buffer (25 mM ammonium bicarbonate and 1.3 mM calcium chloride). The epidermis was homogenised in 8 M urea buffer with an ultrasonicator (S220X, Covaris, Brighton, UK) with 175 W peak power for 8 min. Samples were centrifuged and supernatants diluted to 2 M urea with AB buffer. Dermis- and epidermis-derived supernatants were digested with trypsin SMART Digest Beads (Thermo Fisher Scientific, MA, USA) and agitated overnight at 37°C. Tryptic peptide samples were reduced in 10 mM DTT for 10 min at 60°C, alkylated in 30 mM iodoacetamide for 30 min and acidified in 2% (v/v) trifluoroacetic acid. Biphasic extraction was performed via agitation with ethyl acetate. Peptides were then desalted using OLIGO R3 Reversed-Phase Resin beads (Thermo) and vacuum dried. Samples were re-suspended in 5% acetonitrile (ACN) and peptides (∼10 μg injections) were analysed using LC-MS/MS.

### Mass spectrometry

Dermal- and epidermal-derived peptide mixtures were separately analysed by LC-MS/MS using an UltiMate^®^ 3000 Rapid Separation Liquid Chromatography system (RSLC; Dionex Corporation, CA, USA) coupled to a Q Exactive HF mass spectrometer (Thermo). Peptide mixtures were separated using a multistep gradient from 95% A (0.1% formic acid [FA] in water) and 5% B (0.1% FA in ACN) to 7% B at 1 min, 18% B at 58 min, 27% B at 72 min and 60% B at 74 min at 300 nL min-1, using a 250 mm x 75 μm i.d. 1.7 µM CSH C18, analytical column (Waters, Hertfordshire, UK). Peptides were selected for fragmentation automatically by data-dependant analysis.

### Peptide list generation for peptide location fingerprinting

Raw MS spectrum files were converted to Mascot MGF files containing peak lists with associated mass and intensity values using the ExtractMSn algorithm (Thermo) under default parameters. The Mascot Daemon application v 2.5.1 (Matrix Science; London, UK) was used to automate the submission of peak list MGF files to the Mascot server. The Mascot search engine then correlated the peak spectra within each file to the UniProt human database (2018; Swiss-Prot and TreEMBL). Mascot MS/MS ion searches were performed with the following parameters: database – Swissprot_TreEMBL_2018_01; species – *Homo sapiens*; enzyme – trypsin; peptide charge – 2+ and 3+; max missed cleavages – 2; fixed modifications – carbamidomethyl (mass: 57.02 Da; amino acid: C); variable modification – oxidation (mass: 15.99 Da; amino acid: M); peptide tolerance – 10 ppm (monoisotopic); fragment tolerance – 0.02 Da (monoisotopic); instrument – ESI-TRAP. Mascot search results were exported as DAT files for every run performed. Peptide spectrum matches (PSM) were generated using the Scaffold 4 software (Proteome Software; Portland, OR, USA). DAT files were imported into Scaffold 4 and peptide/protein identifications generated automatically using LFDR scoring. Data were filtered to report only peptides exclusive and unique to their matched proteins. Peptide probability was thresholded to 95% minimum giving a low peptide false discovery rate (FDR) of 0.5% for epidermis samples and 0.6% for dermis samples. Peptide FDR was automatically calculated by Scaffold 4 using peptide probabilities assigned by the Trans-Proteomic Pipeline (Sourceforge; Seattle. WA, USA) using the PeptideProphet™ algorithm. Peptide lists used for peptide location fingerprinting were exported from Scaffold 4 (**Tables S1** and **S2**) within CSV files which were then imported into the MPLF webtool.

### Peptide location fingerprinting using the Manchester Peptide Location Fingerprinting webtool

The MPLF webtool is a publicly available (upon publication in a journal) bioinformatics spectral counting LC-MS/MS analysis tool developed in-house, hosted on our previously published database: the Manchester Proteome website (70). MPLF allows users to perform high-throughput peptide location fingerprinting analysis of LC-MS/MS peptide spectrum lists. Calculations are performed by MPLF on the server side with a code written in Python version 3.8. The back end is powered by Python Django 3.8 framework and by mySQL 8 database and the front end operates REACTjs with Redux state management to ensure speed and reliability of analysis.

The concept of peptide location fingerprinting by the MPLF webtool was designed using LC-MS/MS spectral counting data instead of peak area ion intensity, since spectral counting is the easiest data to access in the field and instrument independent. Additionally, peptide location fingerprinting measures gross differences across the protein structure profile making spectral counting suitable. Lastly, we wanted to make MPLF a useful LC-MS/MS analysis and visualisation tool for other researchers in the field and spectral counting data is easiest for users to access. We hope to continue developing the MPLF webtool and adjust in future to include peak area ion intensity and additional organism proteomes.

The MPLF webtool allows protein amino acid sequence structures to be divided either into 20, 40, 50, 80, 100 aa regional steps or into step sizes corresponding to domains, repeats or regions pre-defined by the UniProt database. Peptide list CSVs (**Tables S1** and **S2**) were imported into the MPLF webtool and peptide spectrum matches (PSM and associated spectral counts) were then automatically summed per respective regional step within a protein, normalised based on individual protein total spectrum count, averaged per group (forearm or buttock; N = 7) and subsequently heat mapped onto representative “amino acid-length scale” schematics of each protein. Average PSM counts corresponding to amino acid number step sizes within protein structures of one group (buttock) were then subtracted from the counts of their corresponding step sizes in the other group (forearm) and divided by each step’s amino acid length to show regional differences in peptide yield. In addition, PSM counts corresponding to each amino acid step size within a protein of one group (buttock) was statistically compared with counts of each corresponding amino acid step size in the other group (forearm) using Bonferroni-corrected, repeated measures paired ANOVA. For further information on peptide location fingerprinting, please refer to previous publications detailing the same process for fibrillin-1 and collagen 6 alpha-3 proteins (10, 19).

### Relative quantification of protein abundance with peak area ion intensity using Progenesis QI software

Relative quantification of protein abundance was performed using Progenesis QI software package (Nonlinear Dynamics, Waters, Newcastle, UK). Raw mass spectra files were imported and ion intensity maps were automatically generated. Ion outlines were automatically aligned by Progenesis QI to a single reference run using default settings. Ion peaks and their relative abundances were then

automatically picked without filtering and normalised to a single reference run by Progenesis QI using default settings. Data were then exported and searched using Mascot v2.5.1 with same search parameters and on the same database as described for peptide location fingerprinting. This was then re-imported back into Progenesis QI where identified peptide ions were automatically matched. Normalised abundance for each protein was calculated by Progenesis QI as the sum of the each matched peptide ion abundance (individual peptide ion abundance is equal to the sum of the intensities within the isotope areas). Normalised protein abundances, compared between matched forearm and buttock samples, were statistically analysed within Progenesis QI using a paired (repeated measured) ANOVA test. A fold change for each protein was also calculated automatically (fold change is defined as the higher average normalised abundance of one group divided by the average normalised abundance of the second group). Principal component analysis (PCA) was performed on the quantified proteins using Python Sklearn package (**Fig. S8**).

### TIMP3 and RPL36 Western blotting

Samples were ran on 4 – 12% NuPAGE gels (Life Technologies, Warrington, UK) and transferred onto Immun-Blot PVDF membranes (Bio-Rad, Watford, UK) using a Trans-Blot Turbo Transfer system (Bio-Rad). Membranes were blocked with 5% (v/v) milk in TBS-T (50 mM Tris-HCl, 5% [v/v] TWEEN 20) for one hour at room temperature and split into two halves along the direction of electrophoresis at ∼40 kDa. Bottom halves of dermal sample membranes were incubated with rabbit anti-TIMP3 (Cell Signalling Technology; D74B10) and top halves with mouse anti-HSA (Abcam, ab10241; loading control). Bottom half of epidermal sample membranes were incubated with rabbit anti-RPL36 (Sigma; HPA047153) and top halves with rabbit anti-VCL (Sigma; HPA063777; loading control). All primary antibodies were incubated at 1:1000 dilution, overnight at 4°C. Membranes were washed thrice with TBS-T before incubation for one hour at room temperature with secondary antibodies (horse radish peroxidase goat anti-mouse or anti-rabbit; Bio-Rad), all at 1:5000 dilution. Blots were developed with Western Lightning Plus ECL (PerkinElmer, Beaconsfield, UK).

### Expansion of the Manchester Proteome

Skin proteins identified (Scaffold 4; peptide probability threshold = 50%, protein probability threshold = 99%, minimum two peptides per protein) in the LC-MS/MS data generated in this study were used to expand the existing Manchester Proteome database with new protein entries (70). All protein identifications with their associated p-values and fold changes are freely searchable on www.manchesterproteome.manchester.ac.uk/proteome.

## Author Contributions

MO and AE contributed equally to this work. MO and AE conceived, designed and performed all proteomic experiments, analysed all the data and prepared the figures. MO programmed and developed the MPLF webtool. AE conceptualised the application of peptide location fingerprinting approach, wrote the paper and performed all western blot experiments. MJS contributed to the conception and design of the study, the interpretation of results, the preparation of figures and to writing. KTM performed all histology, Wiegert’s staining and microscopy of skin tissue. VM and JS contributed to the design of LC-MS/MS sample preparation. SW, ROC and DK provided technical assistance and support for all LC-MS/MS performed and to the interpretation of MS data and figures. CEMG and REBW and contributed to the interpretation of results. JS contributed to the interpretation of peptide location fingerprinting data and the conceptualisation of intragroup variation testing. All authors contributed to reviewing and editing of the paper.

## Funding Disclosure

This study was funded by a programme grant from Walgreens Boots Alliance, Nottingham, UK.

## Competing Interests

The authors declare that they have no conflicts of interest with the contents of this article. Walgreens Boots Alliance has approved this manuscript’s submission but exerted no editorial control over the content.

## Supporting information

List of MPLF analysed dermal proteins sig dif regions

Full list of dermal peptides for peptide fingerprinting

Full list of epidermal peptides for peptide fingerprinting

Intragroup variation testing of dermal protein regions

Intragroup variation testing of epidermal protein regions

List of MPLF analysed epidermal protein sig dif regions

Relative quant of dermis with significant biomarkers

Relative quant of epidermis with significant biomarkers

Dermal biomarker IDs in MPLF vs relative protein quant

Epidermal biomarker IDs in MPLF vs relative protein quant

List of MPLF analysed tendon proteins sig dif regions

## Acknowledgements

We would like to thank all the participants who donated skin for this study and research nurses J. Bastrilles and G. Aarons at the The Dermatopharmacology Unit of Salford Royal NHS Foundation Trust. We would also like to acknowledge the help of the University of Manchester research IT team members, in particular A. Gilchrist for setting up the servers to host the MPLF webtool and J. Selley of the BioMS facility for his aid in bioinformatic analysis during the Covid-19 pandemic.

---

**Table S1.** Full list of LC-MS/MS-identified exclusive peptides in dermis samples used for structural mapping and fingerprinting.

**Table S2.** Full list of LC-MS/MS-identified exclusive peptides in epidermis samples used for structural mapping and fingerprinting.

**Table S3:** Intragroup variation testing of dermal protein regions

**Table S4:** Intragroup variation testing of epidermal protein regions

**Table S5.** Full list of peptide fingerprinted dermal protein regions with significantly different peptide counts between forearm and buttock

**Table S6.** Full list of peptide fingerprinted epidermal protein regions with significantly different peptide counts between forearm and buttock

**Table S7.** Full list of whole protein relatively quantified dermal proteins by peak ion intensity ranked by significance.

**Table S8.** Full list of whole protein relatively quantified epidermal proteins by peak ion intensity ranked by significance.

**Table S9:** Dermal protein biomarkers identified exclusively by peptide location fingerprinting, relative protein quantification or by both methodologies.

**Table S10:** Epidermal protein biomarkers identified exclusively by peptide location fingerprinting, relative protein quantification or by both methodologies.

**Table S11:** Full list of peptide fingerprinted tendon protein regions with significantly different peptide counts between old and buttock

**Figure S1.**
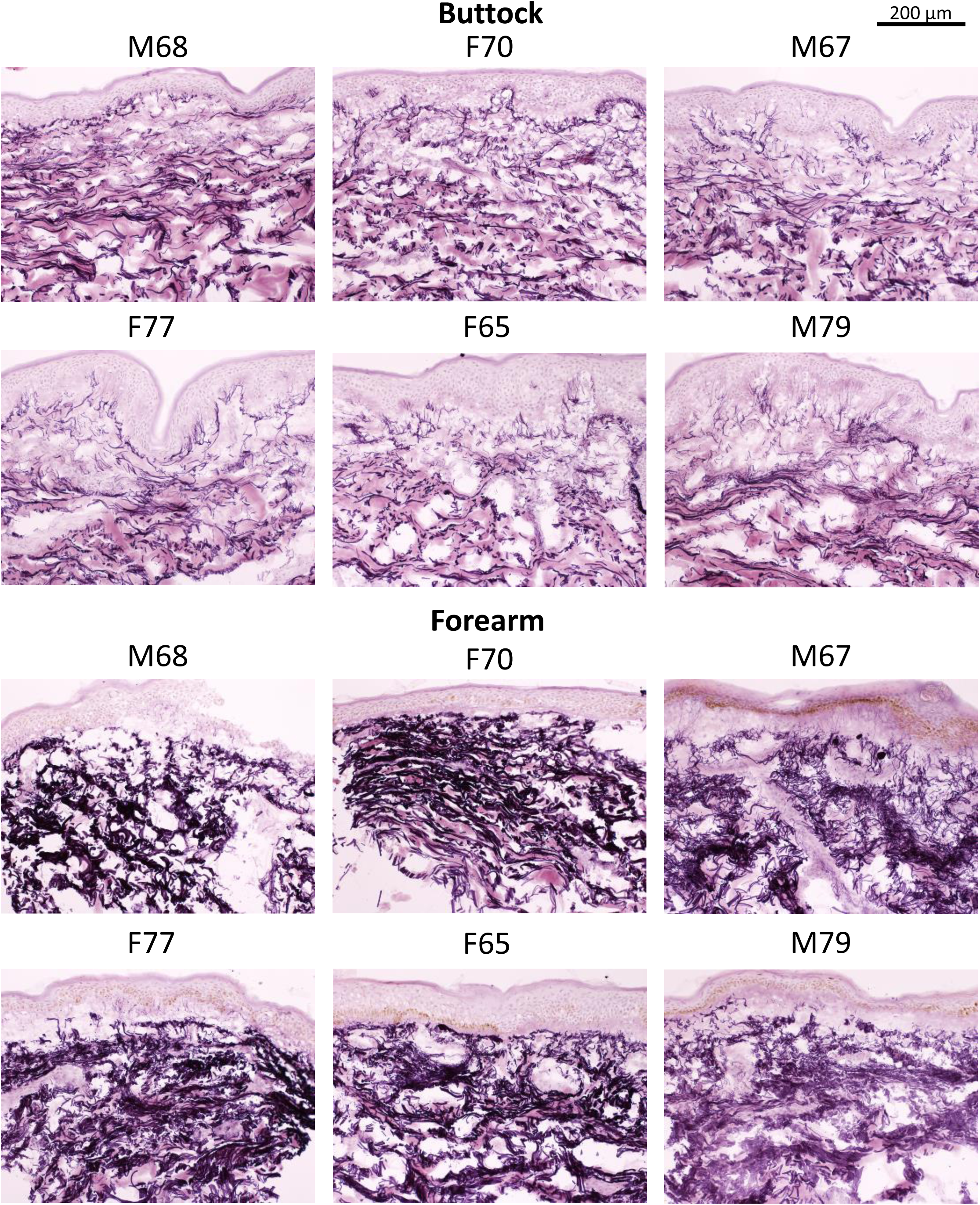
Representative histology images of biopsies to visually showcase photoageing phenotype. .

**Figure S2.**
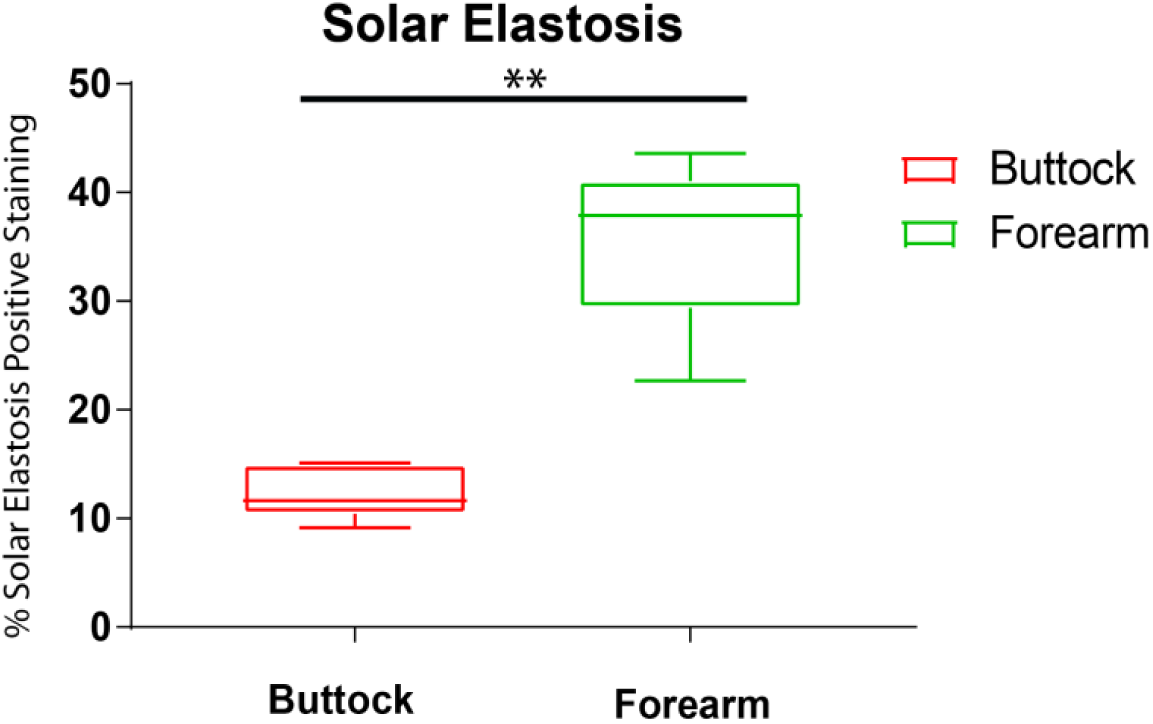
Skin samples from photoexposed forearm had significant solar elastosis compared to photoprotected buttock.

**Figure S3:**
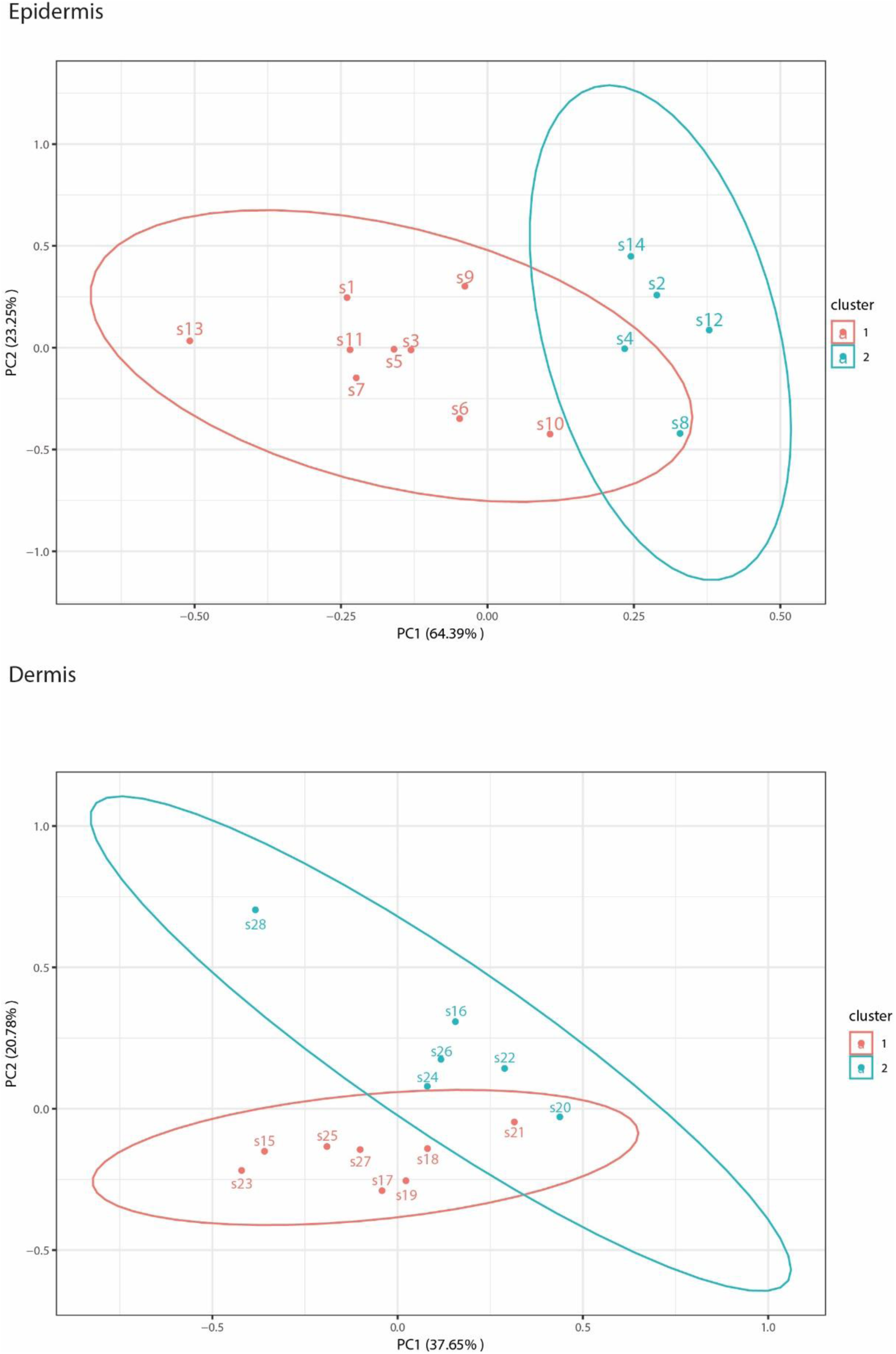
Principal component analysis (PCA) of spectral count data used for peptide location fingerprinting shows clear separation of forearm and buttock data into distinct clusters.

**Figure S4:**
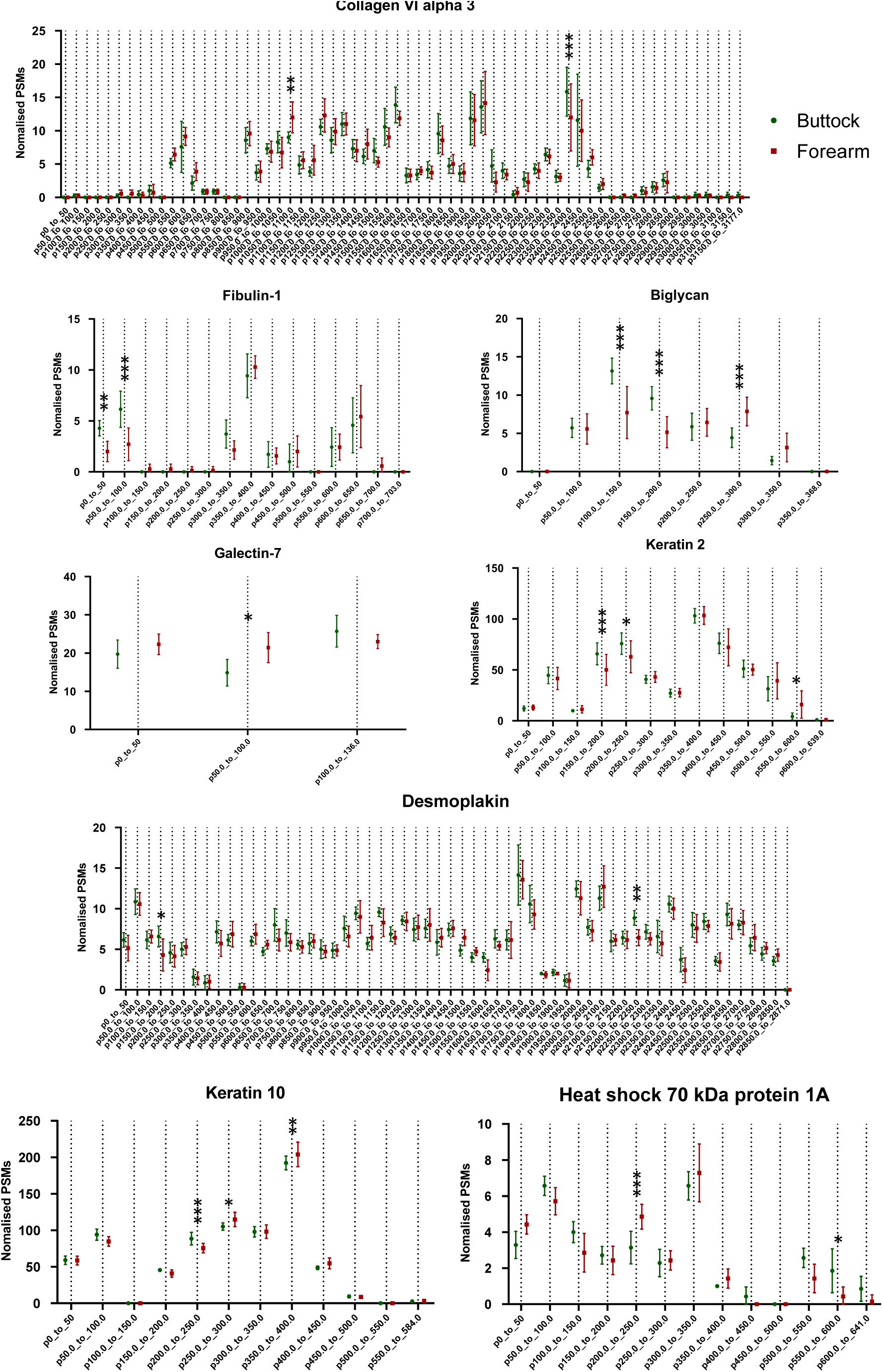
Eight exemplary biomarkers exhibiting photoageing-specific structural modifications.

**Figure S5.**
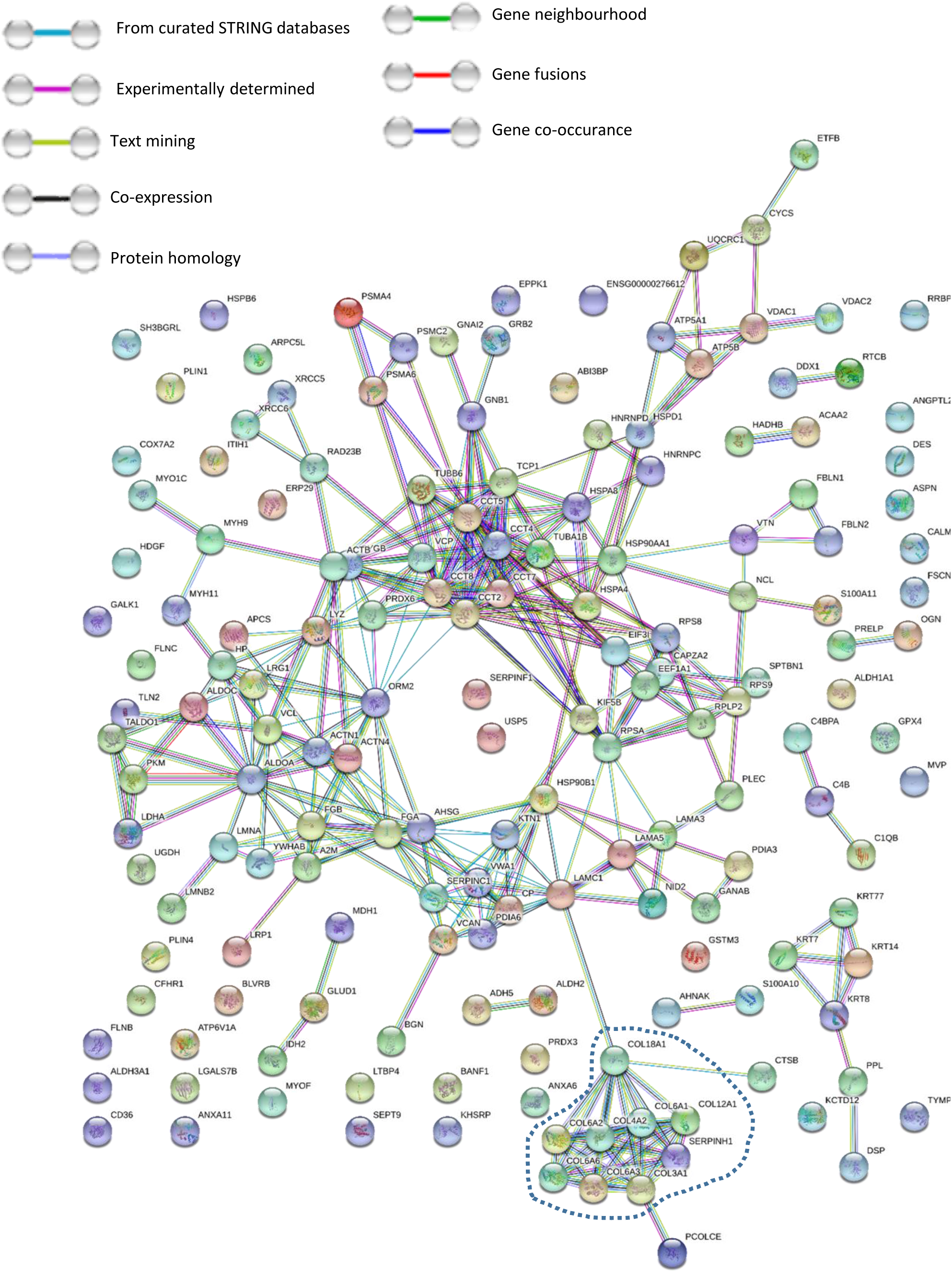
Protein-protein interaction network analysis of dermal biomarkers containing structural modifications indicates a global effect to tissue homeostasis as a consequence of chronic sun exposure.

**Figure S6.**
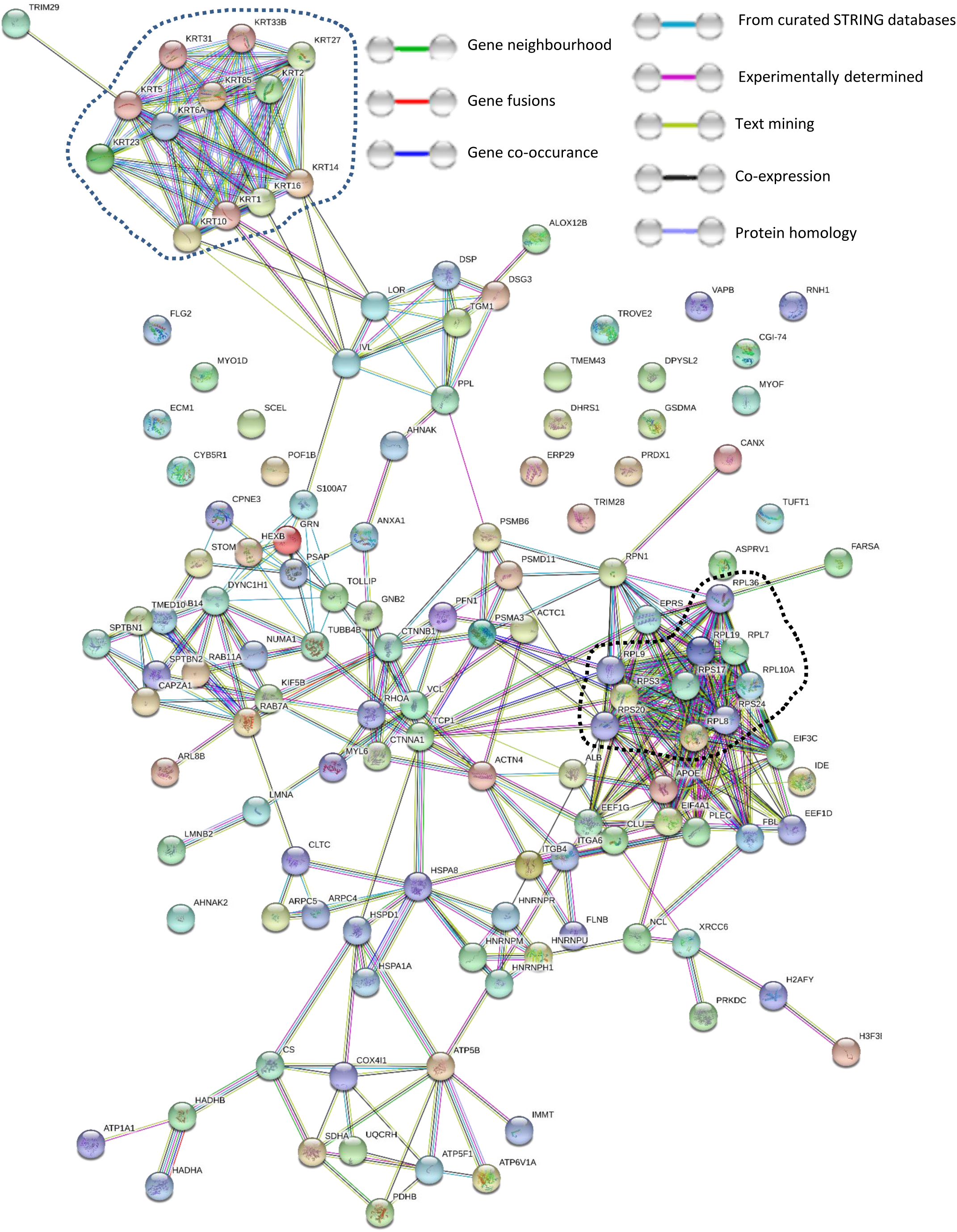
Protein-protein interaction network analysis of epidermal biomarkers containing structural modifications indicates a global effect to tissue homeostasis as a consequence of photoageing.

**Figure S7.**
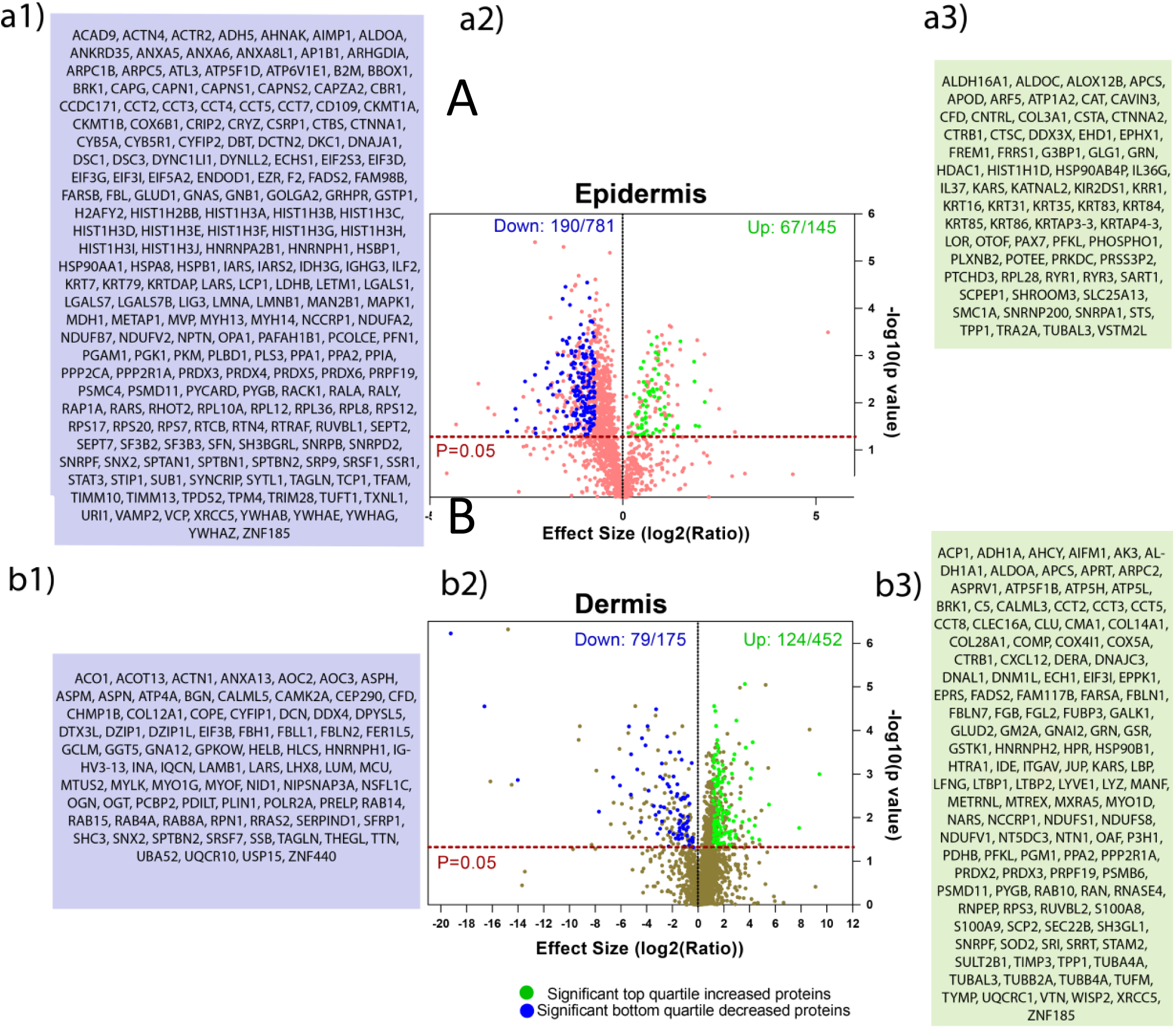
Label-free relative quantification of protein abundance by peak area ion intensity identifies multiple proteins with significant differences in relative abundance between matched photoaged forearm and intrinsically-aged buttock skin.

**Figure S8.**
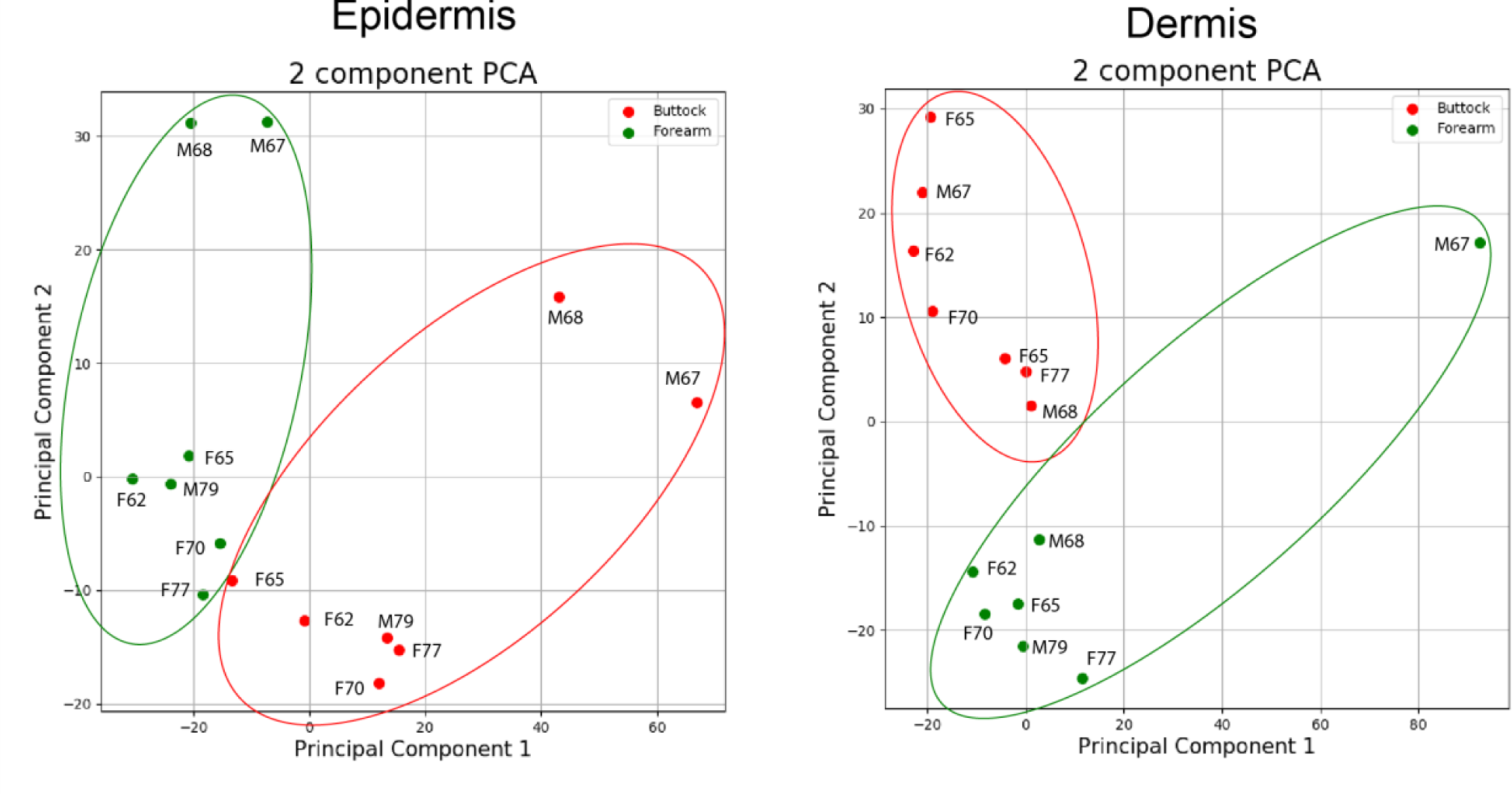
PCA analysis of peak area ion intensity data used for relative quantification shows clear data separation between forearm and buttock samples analysed.

**Figure S9:**
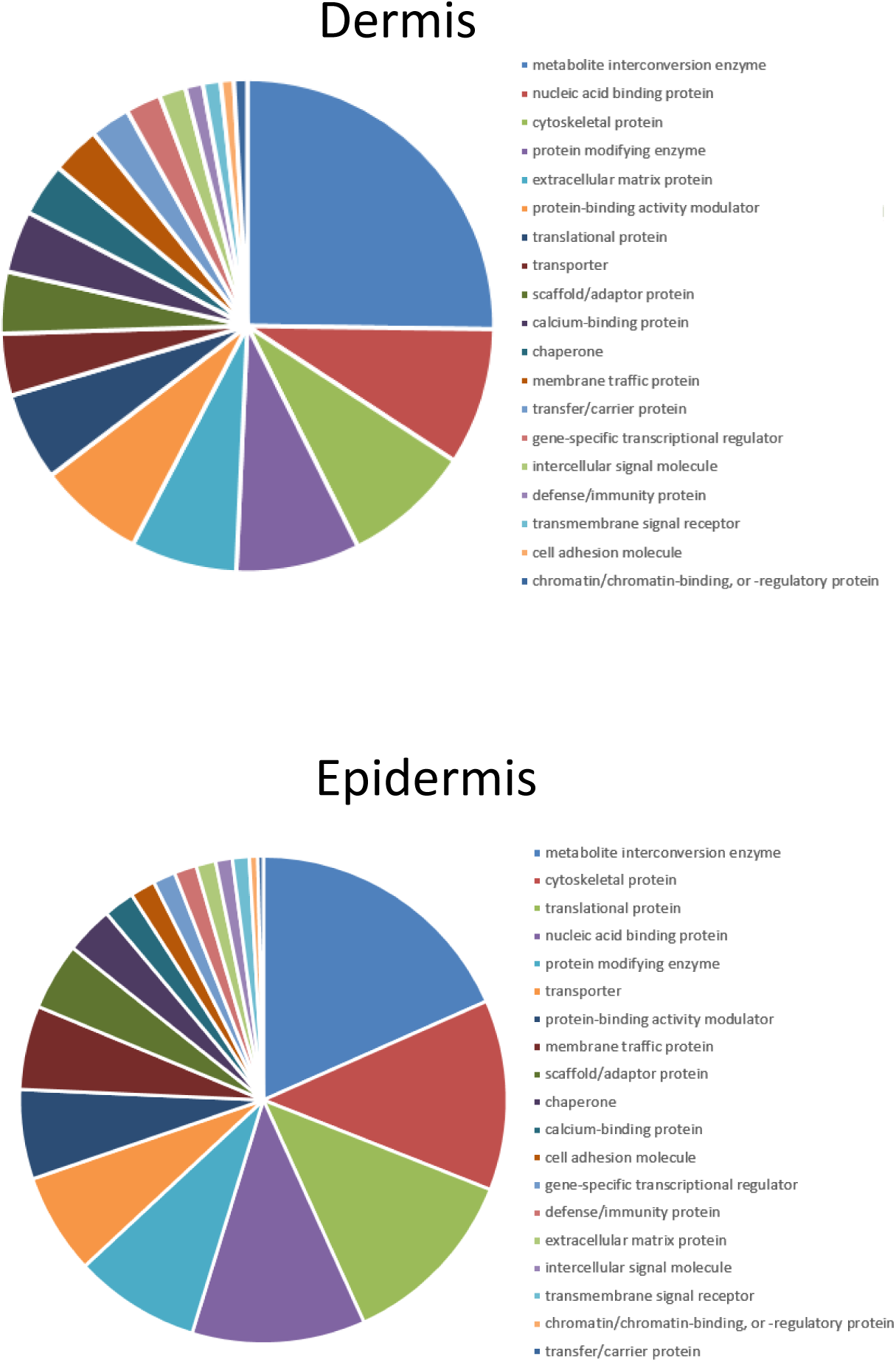
Classification of protein biomarkers significantly different in relative abundance into functional groups reveals metabolite interconversion enzymes, nucleic-acid binding proteins, cytoskeletal proteins and translational proteins as the main classes in skin most affected by the photoageing process.

**Figure S10:**
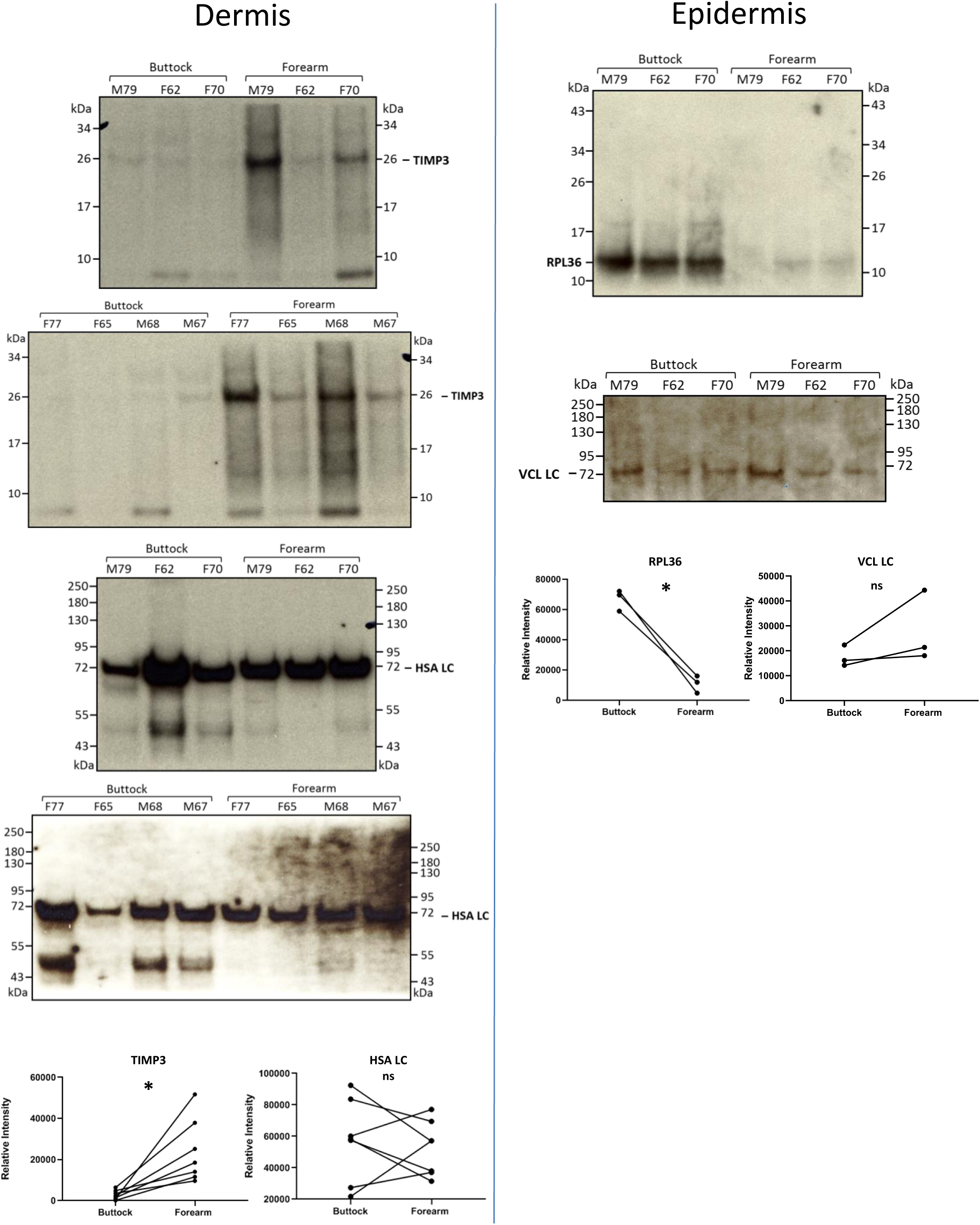
Western blot validation of LC-MS/MS relative quantification of protein abundance highlights TIMP3 and RPL36 as novel biomarkers of photoageing.

**Figure S11:**
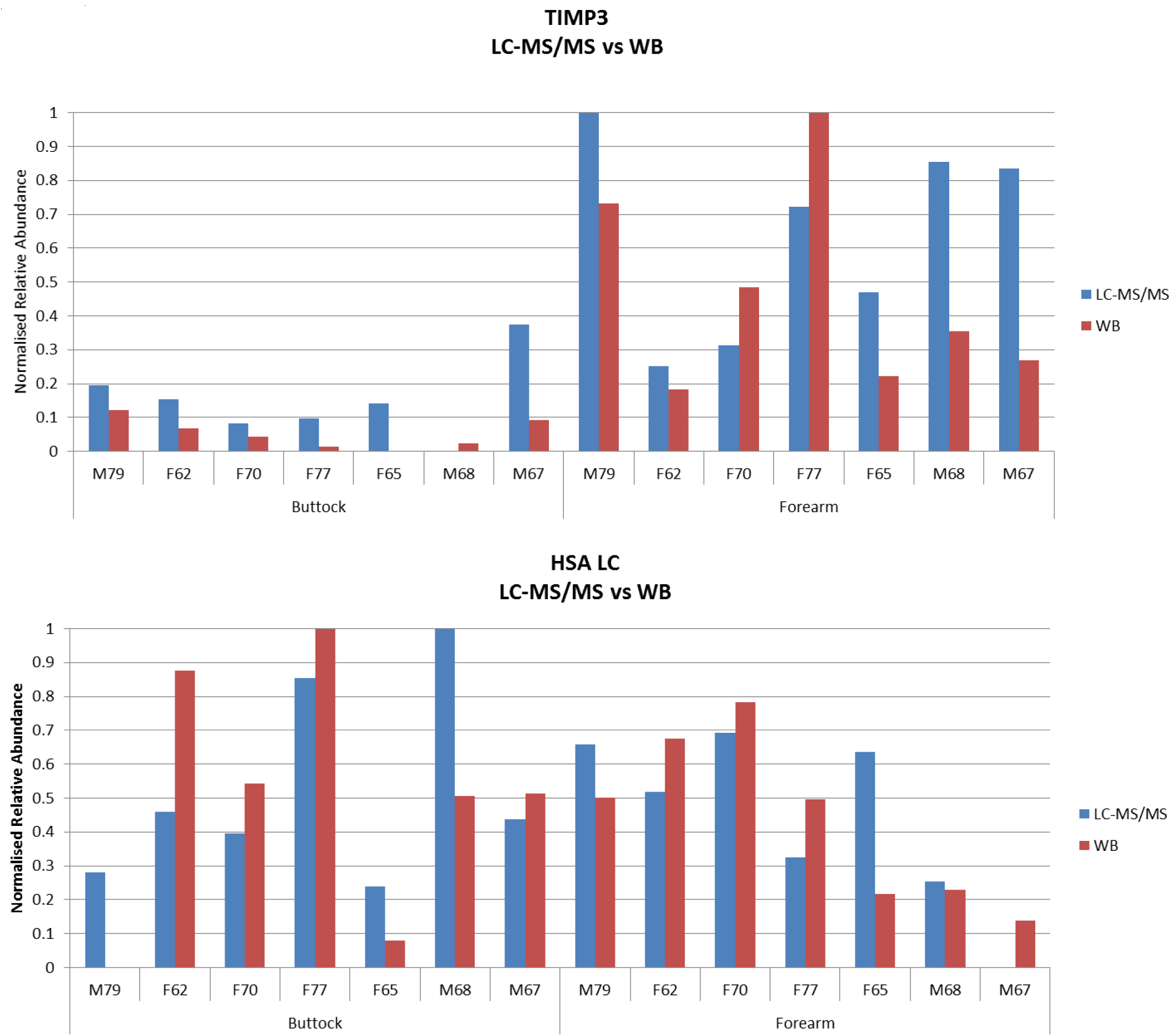
Dermis LC-MS/MS-based relative abundances for TIMP3 and HSA match well with Western blot relative abundances on a sample by sample basis.

## Notes

### Competing Interest Statement

The authors have declared no competing interest.

