## Supplementary material for "Peptide location fingerprinting reveals modification-associated biomarkers of ageing in human tissue proteomes": Figure and Supporting Figure Legends

**Figure S1. Representative histology images of biopsies to visually showcase photoageing phenotype.** Weigert's stained cryosections from aged outer forearm skin samples exhibited marked elastosis and disrupted elastic fibre architecture compared to matched buttock samples (sex and age displayed above images).

**Figure S2. Skin samples from photoexposed forearm had significant solar elastosis compared**

**to photoprotected buttock.** Elastic fibre abundance in forearm skin sections was quantified and compared to that of photoprotected buttock (N = 6, average age = 71 years, range = 65 – 79 years, 3 males; data = median, IQR and range). Elastic fibre abundances in photoexposed forearm skin sections were significantly higher than in photoprotected buttock ($p = 0.0012$, paired t test).

**Figure S3. Principal component analysis (PCA) of spectral count data used for peptide location fingerprinting shows clear separation of forearm and buttock data into distinct clusters.** Odd numbered samples corresponded to buttock, even numbered samples corresponded to forearm. Partitioning around medoids clustering analysis automatically separated buttock (red) and forearm (blue) into two distinct clusters.

**Figure S4. Eight exemplary biomarkers exhibiting photoageing-specific structural modifications.** Interleaved graph representations of proteins found in **Fig. 3** of main text. Proteins were segmented into 50 amino acid-sized step regions with average peptide counts (PSMs; N = 7) statistically compared between forearm and buttock (graphs = average PSMs, error bars = SD; * ≤ 0.05, ** ≤ 0.01, *** ≤ 0.001, Bonferroni-corrected repeated measures paired ANOVA). Regional significant differences in peptide yield along these protein structures can be seen between photoaged forearm and intrinsically aged buttock.

**Figure S5. Protein-protein interaction network analysis of dermal biomarkers containing structural modifications indicates a global effect to tissue homeostasis as a consequence of chronic sun exposure.** STRING analysis (minimum required interaction score = 0.700) was used to look as protein-protein interactions. Clear cluster of interactions can be seen for collagens in particular (dashed blue line).

**Figure S6. Protein-protein interaction network analysis of epidermal biomarkers containing structural modifications indicates a global effect to tissue homeostasis as a consequence of photoageing.** STRING analysis (minimum required interaction score = 0.700) was used to look as protein-protein interactions. Clear clusters of interactions can be seen for keratins (dashed blue line) and ribosomal proteins (dashed black line) in particular.

**Figure S7. Label-free relative quantification of protein abundance by peak area ion intensity identifies multiple proteins with significant differences in relative abundance between matched photoaged forearm and intrinsically aged buttock skin.** Photoageing had a significant effect on protein abundance in human skin. Proteins with relative abundances which were significantly lower in forearm than in buttock (Progenesis QI multivariate paired ANOVA; $p < 0.05$) and in the bottom quartile for fold change (blue points) are listed to the left of the volcano plots. For epidermis, these include histones, chaperones, heat shock proteins, ribosomal proteins, galectins and redox enzymes. For dermis these include proteoglycans and basement membrane proteins. Proteins with relative abundances which were significantly higher in forearm than in buttock and in the top quartile for fold change (green points) are listed to the right of the volcano plots. For epidermis, this includes a number of keratins and, for dermis, a number of collagens, protease modulators and elastic fibre proteins.

**Figure S8. PCA analysis of peak area ion intensity data used for relative quantification shows clear data separation between forearm and buttock samples analysed.** Partitioning around medoids clustering analysis automatically separated buttock (red) and forearm (green) into two distinct clusters.

**Figure S9. Classification of protein biomarkers significantly different in relative abundance into functional groups reveals metabolite interconversion enzymes, nucleic-acid binding proteins, cytoskeletal proteins and translational proteins as the main classes in skin most affected by the photoageing process.** Biomarker proteins with significant differences in abundance between forearm and buttock according to peak area ion intensity analysis were categorised into protein classes (PANTHER classification system; large multi-coloured pie charts: clockwise rankings with top rank at 12:00; only classes with two or more proteins are represented). Metabolite interconversion enzymes, nucleic acid-binding proteins and cytoskeletal proteins were in the top four classes affected for both dermis and epidermis. Protein modifying enzymes for dermis and translational proteins for epidermis were also in the top four for each respective sub tissue. Although the majority of the dermis is comprised of ECM, ECM proteins were not in the top four classes affected. This indicates that relative protein abundance measurements are not as suitable as protein modification measurements by peptide location fingerprinting (**Fig. 4** in main text) at distinguishing photoageing-related differences to ECM proteins.

**Figure S10. Western blot validation of LC-MS/MS relative quantification of protein abundance highlights TIMP3 and RPL36 as novel biomarkers of photoageing.** Relative abundance measurements of two proteins identified as significantly different between matched photoaged forearm and intrinsically aged buttock samples by peak area ion intensity was confirmed by Western blotting. TIMP3 detection was negligible in buttock dermis samples from aged individuals (sex and age in years labelled per lane) but highly present in matched forearm dermis samples (loading control = human serum albumin – HSA). Quantification of relative intensity shows that TIMP3 abundance is significantly higher in forearm than in matched buttock (paired t test, p = 0.0115). In contrast, RPL36 was highly present in buttock epidermis samples from aged individuals but negligibly detected in matched forearm epidermis samples (loading control = vinculin, VCL). Quantification of relative intensity shows that RPL36 abundance is significantly lower in forearm than in matched buttock (paired t test, p = 0.0112). Loading controls (HSA for dermis samples and VCL for epidermis samples) were chosen based on their lack of significant differences in relative abundance between forearm and buttock samples according to LC-MS/MS peak area ion intensity. Western blot analysis of these controls showed no significant differences between forearm and buttock samples also corroborates peak area intensity comparisons.

**Figure S11. Dermis LC-MS/MS-based relative abundances for TIMP3 and HSA match well with Western blot relative abundances on a sample by sample basis.** TIMP3 and HSA relative abundance values for LC-MS/MS peak area ion intensity were normalised against relative abundance value for Western blotting band intensities to be compared on a sample by sample basis (1 = highest relative abundance for each method and anything below is represented as a fraction of that highest value).
